# Deciphering Tumor Microenvironment Dynamics in Tumorigenesis and Lymph Node Metastasis of Esophageal Squamous Cell Carcinoma using Single-cell RNA Sequencing

**DOI:** 10.1101/2025.11.27.690370

**Authors:** Hansoll Na, Tae Hee Hong, Chung Lee, Young Ho Yang, Ha Eun Kim, Byung Jo Park, Hyun Ki Kim, Dae Joon Kim

## Abstract

Esophageal squamous cell carcinoma (ESCC) exhibits profound inter-tissue heterogeneity, yet how immune and stromal ecosystems diverge between primary tumors and metastatic lymph nodes remains poorly understood. Here, we generated a single-cell atlas of 344,790 cells and paired T cell receptor profiles from primary tumor mucosa, adjacent mucosa, non-metastatic lymph nodes, and metastatic lymph nodes from 18 patients with treatment-naive ESCC. We identify compartment-specific immune and stromal architectures that shape distinct antitumor responses and suppressive programs. Primary tumors were characterized by marked enrichment of regulatory T cells (T_REG_) and activated dendritic cells coordinated through CTLA4-associated circuits, accompanied by accelerated CD8^+^ T cell differentiation toward intermediate and terminal exhaustion with loss of cytotoxicity. In contrast, metastatic lymph nodes preserved substantial pools of pre-exhausted CD8^+^ T cells with retained effector potential and strong clonal connectivity to the primary tumor, suggesting sustained antigen-driven trafficking. However, these reinvigoration-competent populations were embedded within a niche dominated by *TREM2^high^* macrophages that delivered SPP1-dependent suppressive signals to exhausted and regulatory T cells. Stromal lineages also displayed niche-specific specialization: tumor mucosa contained chemokine-rich inflammatory fibroblasts, whereas metastatic lymph nodes upregulated extracellular matrix remodeling programs. Together, our findings demonstrate that ESCC progression is governed by anatomically distinct exhaustion trajectories, clonal behaviors, stromal states, and suppressive circuits. This framework provides mechanistic insight into why lymph node response is a key determinant of clinical outcome and highlights the need for site-tailored immunomodulation strategies that target T_REG_-mediated suppression in primary tumors and *TREM2^high^* macrophage programs in metastatic nodes.

## Introduction

Esophageal squamous cell carcinoma (ESCC) represents a substantial global health burden as the eleventh most common cancer, and the seventh leading cause of cancer related mortality [1]. In East Asia, including South Korea, over ninety percent of esophageal cancer cases are squamous cell carcinoma, underscoring its regional and clinical relevance [2, 3]. Despite advances in diagnostic imaging, surgical techniques, and chemoradiotherapy, ESCC remains a highly challenging malignancy with persistently poor prognosis and limited therapeutic options [4]. In South Korea, the five-year survival rate remains under 40%, highlighting the urgent need for improved biomarkers and therapeutic strategies [5, 6].

Single-cell RNA sequencing has transformed our understanding of tumor ecosystems by resolving cellular heterogeneity, defining rare or transitional states, and characterizing interactions within the tumor microenvironment [7]. While single cell studies have begun to elucidate features of primary ESCC [8, 9], the molecular determinants of metastatic progression remain incompletely defined. In particular, how immune and stromal landscapes remodel as tumor cells disseminate to lymph nodes is not well understood.

Recent clinical studies have suggested that although neoadjuvant chemoradiotherapy induces strong responses in primary tumors, the addition of immune checkpoint blockade enhances lymph node control and improves both overall and disease-free survival [10]. These observations indicate that lymph node burden is a critical determinant of outcome in ESCC and further imply that immune regulation in lymphoid niches differs from that in the primary tumor. However, from the standpoint of cellular composition, functional state, and lineage specific remodeling, it is unclear whether preoperative immune checkpoint blockade more effectively modulates the lymph node environment than the tumor itself.

To address these gaps, we performed single-cell RNA sequencing of paired primary tumor mucosa, adjacent mucosa, non-metastatic lymph nodes, and tumor associated metastatic lymph nodes from eighteen ESCC patients. By comparing these distinct anatomical niches, we aimed to define the immune and stromal architectures that accompany metastatic dissemination and to identify T cell subsets with potential for reinvigoration. Our findings reveal compartment specific microenvironments and highlight unique immunosuppressive circuits in metastatic lymph nodes. Moreover, we characterize CD8^+^ T cell states with evidence of plasticity, suggesting opportunities to enhance immunotherapeutic responses in ESCC.

## Results

### The immune microenvironment of esophageal squamous cell carcinoma is compartmentalized across tumor and lymphoid niches

To generate a comprehensive cellular atlas of ESCC, we performed single-cell RNA sequencing on 58 specimens from 18 treatment naive, resectable patients. Samples included primary tumor mucosa (TM), adjacent normal mucosa (NM), normal lymph nodes (NLN), and tumor-associated metastatic lymph nodes (TLN) (Figure 1a). Clinicopathological characteristics of the cohort are summarized in Table S1.

**Figure. 1.**
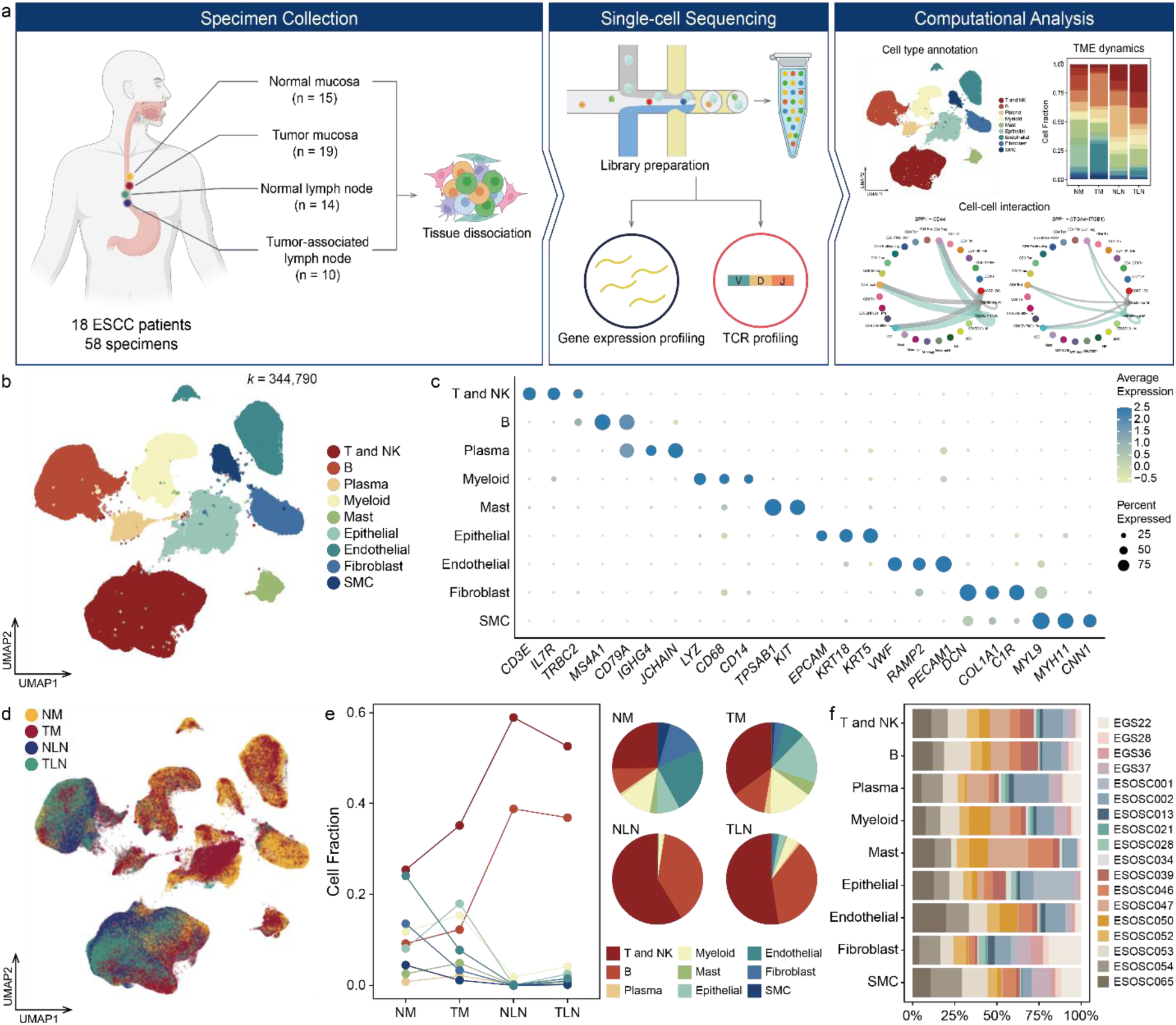
Cellular landscape of ESCC microenvironment. (a) Overview of the study workflow. Fresh tissues were collected from normal mucosa (NM), tumor mucosa (TM), normal lymph nodes (NLN), and tumor-associated lymph nodes (TLN) from 18 patients with esophageal squamous cell carcinoma (ESCC) and processed for single-cell RNA sequencing and T cell receptor (TCR) profiling. (b) Uniform manifold approximation and projection (UMAP) of 344,790 single-cell transcriptomes, annotated into nine major cellular lineages: T and NK cells, B cells, plasma cells, myeloid cells, mast cells, epithelial cells, endothelial cells, fibroblasts, and smooth muscle cells (SMC). (c) Dot plot illustrating lineage-specific marker genes, confirming robust cell-type identification. Dot size indicates the proportion of cells expressing the marker, while the color intensity reflects the normalized expression level. (d) UMAP visualization colored by tissue origin. (e) Relative abundance of each lineage across the four tissue niches. Line plot depicting dynamic changes, and pie charts summarize the overall cellular composition within each microenvironment. (f) Distribution of annotated lineages across individual patients.

After quality filtering, doublet removal, reference-based confidence filtering using Azimuth [11], a total of 344,790 high quality cells were retained with an average of 1,888 detected genes and 5,704 UMIs per cell (Figure S1a). The final dataset consisted of 78,289 NM cells, 104,523 TM cells, 111,082 NLN cells, and 50,896 TLN cells, providing broad representation across anatomical niches (Figure S1b).

Dimensionality reduction using principal component analysis followed by UMAP embedding identified nine major cellular lineages, including T and NK cells, B cells, plasma cells, myeloid cells, mast cells, epithelial cells, fibroblasts, smooth muscle cells, and endothelial cells (Figure 1b). Annotation was supported by the expression of canonical markers such as *CD3E*, *IL7R* for T cells, *MS4A1*, *CD79A* for B cells, *IGHG4*, *JCHAIN* for plasma cells, *LYZ*, *CD68* for myeloid cells, *TPSAB1*, *KIT* for mast cells, *EPCAM*, *KRT18* for epithelial cells, *VWF*, *PECAM1* for endothelial cells, *DCN*, *COL1A1* for fibroblasts, and *MYL9*, *MYH11* for smooth muscle cells (Figure 1c and Figure S2).

Both pathological lymph node negative (pN0) and positive (pN^+^) patients contributed to each major lineage without disproportionate enrichment, confirming balanced representation of disease stages across cell types (Figure S1c). Aggregating cells across all tissues further showed that T and NK cells were the most abundant lineage in the dataset, followed by myeloid, epithelial, and fibroblast compartments (Figure S1d). With this global distribution established, we next compared how these lineages were allocated across anatomical sites.

Comparison of cellular compositions revealed marked differences between compartments (Figure 1d, e). T and NK cell fractions were increased in TM compared to NM, suggesting immune infiltration into the tumor. Endothelial cells were notably reduced in TM, implying compromised vascular integrity. In lymphoid tissues, immune lineages dominated as expected; however, TLN displayed a significant expansion of epithelial and endothelial populations relative to NLN, reflecting structural remodeling associated with metastatic colonization. These nine cell types were detected in almost every patient, but their proportions varied (Figure 1f).

To further resolve cellular heterogeneity, we performed subclustering within major lineages, identifying 19 T and NK subsets, 10 B and plasma cell subsets, 9 myeloid subsets, and 9 fibroblasts and smooth muscle subsets. These refined annotations provided the foundation for subsequent analyses of lineage specific remodeling across tumor and lymph node microenvironments.

### T cell dynamics reveal regulatory dominance in mucosa and selective effector enrichment in metastatic lymph nodes

T and NK cells, which are central to immunotherapy, constituted the largest immune component in the ESCC ecosystem. To resolve their heterogeneity, we analyzed all 148,526 T and NK cells and performed high resolution subclustering, identifying 18 transcriptionally distinct subgroups (Figure 2a). Annotation was guided by canonical marker expression patterns, as shown in the dot plot summarizing average gene expression and percent expression across subsets (Figure S3a). These subgroups encompassed CD4^+^ and CD8^+^ naïve T cells (T_N_) (*LEF1, SELL*), CD4^+^ memory T cells (T_M_) (*ANXA1*, *LMNA*), CD4^+^ tissue resident memory T cells (T_RM_) (*CXCR6*, *CD69*, *ITGAE*), CD4^+^ regulatory T cells (T_REG_) (*FOXP3*, *CTLA4*), CD4^+^ follicular helper like T cells (T_FH-like_) (*CXCL13*, *SH2D1A*), CD4^+^ stressed T cells (T_STR_) (*HSPA6*, *BAG3*) [12], CD4^+^ *IGF1R^+^* T cells, CD8^+^ effector memory T cells (T_EM_) (*GZMK*, *GZMA*), CD8^+^ terminally differentiated effector memory and effector cells (T_EMRA_) (*CX3CR1*), CD8^+^ *ZNF683^+^* and *ZNF683^-^* T_RM_ populations, CD8^+^ *IFNG-AS1^+^* T cells, CD8^+^ exhausted T cells (T_EXH_) (*PDCD1*, *LAG3*, *HAVCR2*), CD8^+^ proliferating exhausted T cells (*HAVCR2*, *MKI67*, *TOP2A*), *CD117^+^* T cells, mitochondria high T cells (MT-high) (*MT-CO1*, *MT-ND6*), and NK cells (*NCAM1*, *KIR2DL4*).

**Figure. 2.**
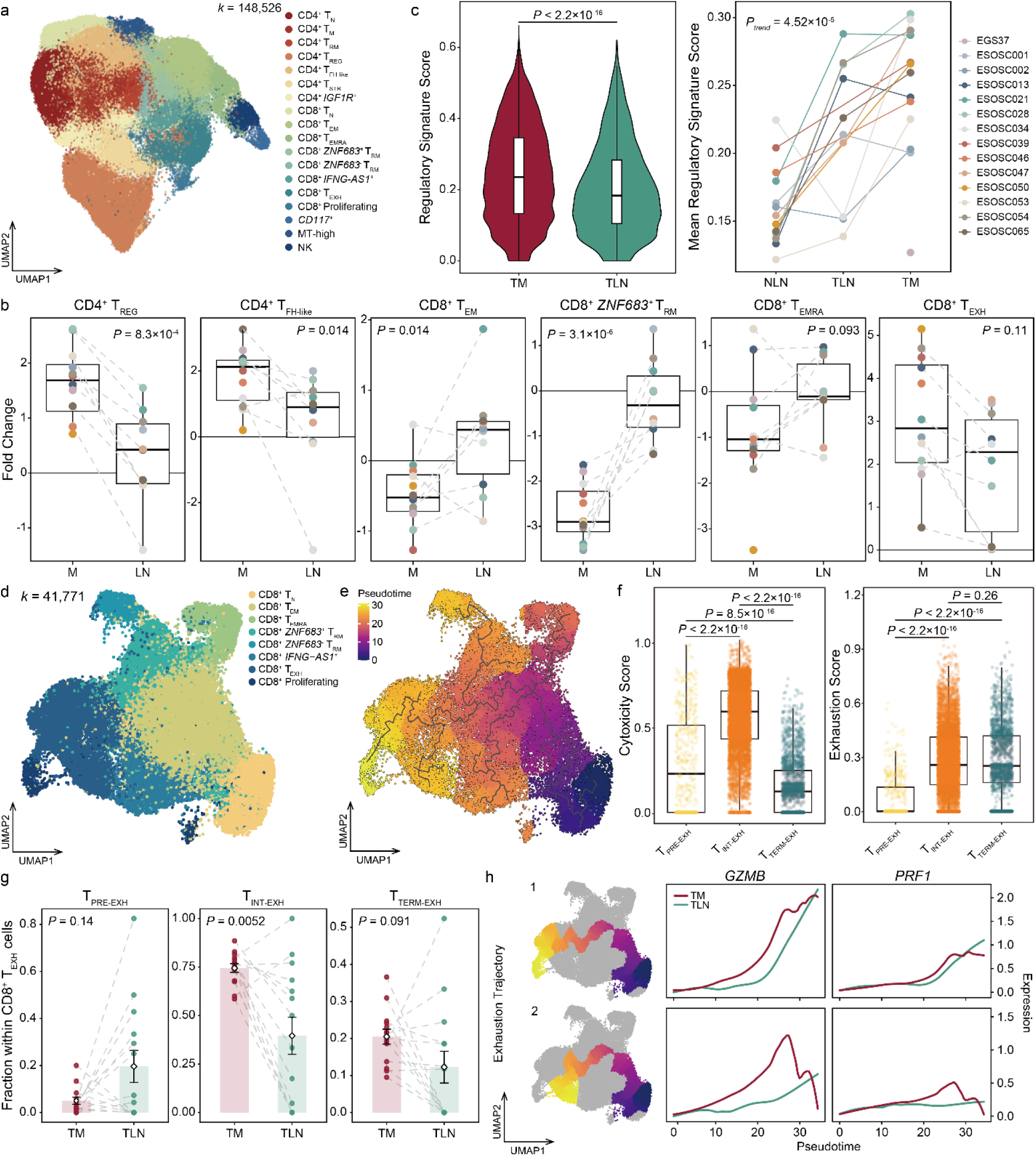
T cell heterogeneity, tissue-specific remodeling, and bifurcated exhaustion trajectories in ESCC. (a) UMAP of 148,526 T and NK cells partitioned into 18 subclusters. (b) Box plots comparing fold changes of fractions of major T cell subsets between TM versus NM and TLN versus NLN. Fold changes were calculated per patient (n = 14), and significance assessed using two-sided Wilcoxon rank-sum test. Data are represented as mean ± S.D. (c) Regulatory signature scores of T_REG_ cells in TM and TLN (two-sided Wilcoxon rank-sum test), and monotonic trends across NLN, TLN (one-sided Jonckheere-Terpstra test). (d) UMAP of 41,771 CD8^+^ T cells. (e) Pseudotime projection of CD8^+^ T cells. (f) Cytotoxicity and exhaustion signature scores across CD8⁺ T_EXH_ cell states (two-sided Wilcoxon rank-sum test). (g) Patient-level fractions of pre-exhausted (T_PRE-EXH_), intermediate (T_INT-EXH_), and terminally exhausted (T_TERM-EXH_) CD8⁺ T cells in TM and TLN (two-sided Wilcoxon rank-sum test). (h) Expression dynamics of cytotoxicity-related genes (*GZMB* and *PRF1*) along the two exhaustion trajectories.

To examine compartment-specific alterations, we focused on pN^+^ patients (n = 131,914 cells). For each patient, fold changes in the abundance of T cell subsets were calculated between TM and NM, and between TLN and NLN. Across samples, major T cell states were comparably represented in each tissue and showed broadly consistent distributions across individuals, providing a balanced foundation for estimating relative changes in abundance (Figure S3b, c). Most subsets showed significant compartment-specific differences, although T_EMRA_ displayed only marginal enrichment in TLN and T_EXH_ expanded in both TM and TLN without reaching statistical significance (Figure 2b). Among the significant shifts, T_REG_ exhibited the most robust and consistent increase, revealing a pronounced enrichment of T_REG_ in primary tumors relative to TLN. Notably, T_REG_ abundance was elevated both quantitatively and qualitatively. Aggregate and per patient tests consistently showed higher regulatory signature scores in TM than TLN, supported by a one-sided Jonckheere–Terpstra trend test (Figure 2c).

T_FH-like_ cells were also relatively enriched in TM. CD4^+^ *CXCL13^+^* T cells have been reported to participate in tertiary lymphoid structure (TLS) formation and B cell–helping niches in several solid tumors, including breast [13] and ovarian cancer [14]. The elevated follicular-helper signature we observed in TM (Figure S3d) suggests that ESCC mucosa similarly provides a microenvironment permissive to the accumulation of T_FH-like_ cells, with functional coordination with B cell programs examined in later sections.

In contrast, cytotoxic effector subsets including CD8^+^ T_EM_ and *ZNF683^+^* T_RM_ were more abundant in TLN than in TM. These patterns suggest that metastatic lymph nodes preserve a more immune active environment enriched for effector and cytotoxic phenotypes, whereas primary tumors are dominated by immunosuppressive circuits driven by T_REG_. The divergence in T cell programs across mucosal and lymphoid niches may contribute to site-specific differences in immunotherapeutic responses. Given these compartment-specific distributions, we next examined the differentiation continuum and activation potential of CD8^+^ T cell subsets through pseudotime and clonal analyses.

### Bifurcated exhaustion trajectories of tumor-infiltrating CD8^+^ T cells emerge across tumor mucosa and lymph nodes

Given the functional relevance of exhausted CD8^+^ T cells in antitumor immunity and the observed trend toward increased abundance in TM, we next examined CD8^+^ T cell differentiation in greater detail. A total of 41,771 CD8^+^ T cells were subset and projected onto a developmental trajectory (Figure 2d, e). Pseudotime analysis revealed a shared progression from CD8^+^ T_N_ cells toward an effector memory state, followed by a divergence into three terminal fates: T_EMRA_, T_RM_, and T_EXH_ (Figure 2e). Across tissues, CD8^+^ T cells in TM occupied significantly more advanced pseudotime states compared with those in TLN, indicating accelerated differentiation within the tumor bed (Kolmogorov–Smirnov test, *P* < 2.2 × 10⁻¹⁶) (Figure S4a). v

Because exhausted T cells represent a key therapeutic target, we next examined T_EXH_ heterogeneity. Unsupervised reclustering resolved four transcriptionally distinct exhausted T cell subsets that distributed along pseudotime in a continuous yet structured manner (Figure S4b, c). Two subsets (exh-1 and exh-2) occupied early pseudotime positions and retained appreciable cytotoxic potential. We refer to these as pre-exhausted CD8^+^ T cells (T_PRE-EXH_). The remaining two subsets localized to later pseudotime positions but differed markedly in effector function: one subset (exh-3) exhibited the most profound loss of cytotoxic signature while retaining high exhaustion signatures, characteristic of terminal exhaustion (T_TERM-EXH_). In contrast, the other (exh-4) maintained relatively high cytotoxicity despite a similar exhaustion magnitude, consistent with an intermediate exhausted phenotype (T_INT-EXH_) (Figure 2f). Hence, T_EXH_ heterogeneity reflects three functionally distinct exhaustion states—pre-exhausted, intermediate, and terminal—distinguished by coordinated differences in pseudotime position, exhaustion depth, and cytotoxic capacity.

The distribution of exhausted states varied substantially across tissues (Figure 2g). Comparisons within pN^+^ patients showed higher proportions of pre-exhausted cells in TLN (*P* = 0.14), whereas intermediate-exhausted cells and terminally exhausted cells were significantly more frequent in TM (*P* = 0.0052 and *P* = 0.091, respectively). Consistent with their earlier differentiation status, T_PRE-EXH_ cells in TLN expressed markedly higher levels of *TCF7*, a defining marker of progenitor-like exhausted CD8^+^ T cells (Figure S4d). These patterns indicate that CD8^+^ T cells in TLN remain in earlier, more plastic exhaustion states, while those in TM progress toward less reversible intermediate or terminal states. Such differences in developmental plasticity, rather than inhibitory receptor magnitude, likely underlie the improved responsiveness of lymphoid compartments to checkpoint blockade.

Mapping these subsets onto developmental trajectories uncovered a bifurcation in exhaustion pathways (Figure 2h). Along exhaustion trajectory 1, pre-exhausted cells progressed toward T_INT-EXH_ while maintaining robust cytotoxic gene expression, suggesting preservation of effector potential even at advanced pseudotime. Along exhaustion trajectory 2, pre-exhausted cells transitioned toward T_TERM-EXH_, accompanied by a pronounced collapse of cytotoxic programs. This collapse was especially marked in TM, where terminally exhausted cells showed sharp declines in *GZMB* and *PRF1* and expression at late pseudotime. By contrast, cytotoxicity was better preserved along both trajectories in TLN, indicating that the lymph node microenvironment slows or mitigates the terminal dysfunction observed in primary tumors.

Together, these findings reveal that metastatic lymph nodes harbor CD8^+^ T cell populations enriched for earlier, more plastic exhaustion states, whereas the tumor mucosa drives CD8^+^ T cells toward terminal dysfunction with irreversible cytotoxic decline. This divergence provides a mechanistic basis for the enhanced responsiveness of lymphoid compartments to immune checkpoint blockade observed clinically, and highlights pre-exhausted CD8^+^ T cells in TLN as a potential source of revivable antitumor immunity.

### scTCR-seq reveals antigen-driven trafficking and compartment-specific cytotoxic programs in CD8^+^ T cells

To investigate whether CD8^+^ T cell responses were coordinated between primary tumors and metastatic lymph nodes, we performed single-cell TCR sequencing on a sub-cohort of eight ESCC patients among whom paired TM and TLN samples were available for four patients. Clonal overlap analysis revealed that TCR repertoires of TM and TLN were substantially more similar than those of NLN and TLN, with the TM–TLN Szymkiewicz–Simpson overlap coefficient showing a consistent increase across all patients (Figure 3a). Although the difference did not reach conventional statistical significance (*P* = 0.062, one-sided Wilcoxon signed-rank test), the median overlap was 3.25-fold higher for TM–TLN than NLN–TLN, suggesting active antigen-driven trafficking of tumor-responsive T cells from primary to metastatic sites (Figure S5a, b).

**Figure. 3.**
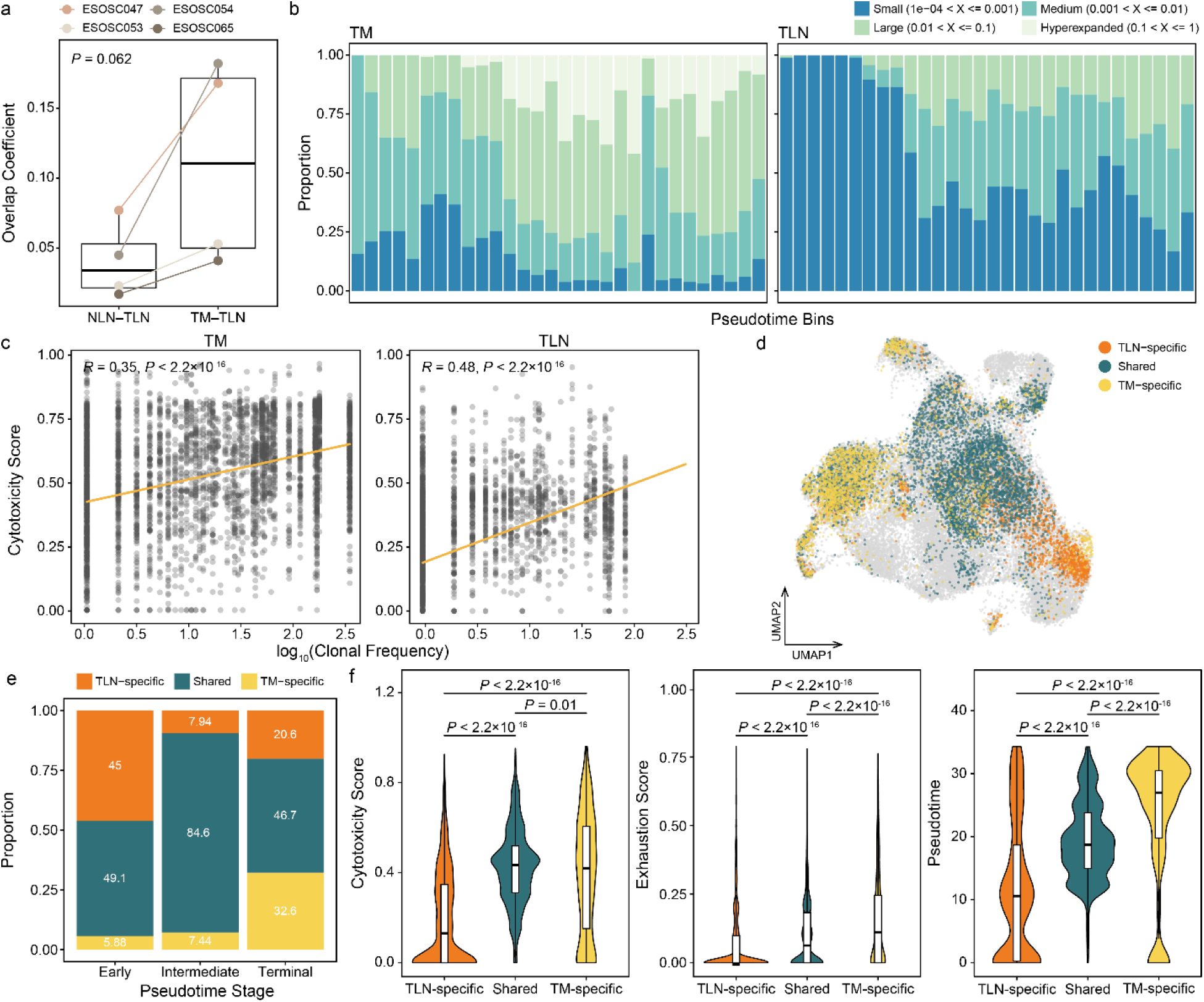
Clonal architecture and functional states of CD8^+^ T cells across tissue niches. (a) Per-patient Szymkiewicz–Simpson overlap coefficients comparing clonal sharing between NLN–TLN and TM–TLN (one-sided Wilcoxon signed-rank test). (b) Distribution of CD8⁺ T cell clone sizes along the exhaustion pseudotime trajectory in TM and TLN. Clonal groups are classified as small, medium, large, or hyperexpanded based on clone size. (c) Scatter plots showing the relationship between clonal frequency and cytotoxicity scores in TM and TLN. Pearson’s correlation coefficient is applied. (d) UMAP of CD8⁺ T cells annotated by clonal tissue specificity, classified as TM-specific, TLN-specific, or shared between both sites. (e) Proportional distribution of TM-specific, TLN-specific, and shared clones across early, intermediate, and terminal pseudotime stages. (f) Cytotoxicity, exhaustion, and pseudotime scores of clone groups stratified by tissue specificity (two-sided Wilcoxon rank-sum test).

To determine how proliferative history shaped CD8^+^ T cell functionality across compartments, we stratified clonal expansions into four size classes: small, medium, large and hyperexpanded. Primary tumors contained markedly larger clones than TLNs, and clone size increased along pseudotime in both tissues (Figure 3b). These trends likely reflect chronic antigenic stimulation in TM, whereas TLNs maintain a more dynamic and less terminally differentiated T cell pool. Cytotoxicity scores correlated positively with clonal expansion in both sites, but the association was stronger in TLN (*R* = 0.48) than TM (*R* = 0.35), indicating that proliferating CD8^+^ T cells in lymph nodes preserve effector potential more effectively than those in TM (Figure 3c). Consistent with this, cytotoxicity scores increased monotonically across clone-size categories, from small to hyperexpanded, in both TLN and TM (Figure S5c), confirming that expanded clonotypes retain robust effector capacity. By contrast, exhaustion scores showed comparable correlations with clonal size in TM and TLN, implying that clonal growth in tumors is more tightly coupled to exhaustion than effector activity (Figure S5d).

We next annotated TCR clonotypes according to their tissue distribution as TM-specific, TLN-specific, or shared across compartments (Figure 3d). Mapping these classifications along pseudotime revealed strikingly distinct developmental trajectories (Figure 3e, Figures S6a). TLN-specific clones were enriched at early pseudotime stages and declined progressively toward later states, consistent with an influx of newly activated cells into lymph nodes. Shared clonotypes, representing cells detectable in both TM and TLN, were most abundant at intermediate stages, whereas TM-specific clones accumulated prominently at late pseudotime. Stage-wise differences in clonotype composition were statistically significant across relevant pairwise comparisons by Fisher’s exact tests after FDR correction (Table S2).

Functional profiling further distinguished the three groups (Figure 3f). TM-specific clones displayed significantly higher pseudotime values, consistent with their greater differentiation and chronic stimulation within tumors. Shared clones, however, retained cytotoxicity comparable to TM-specific clones despite occupying earlier pseudotime states, suggesting that antigen-experienced T cells traffic between TM and TLN while maintaining effector potential. TLN-specific clones showed markedly reduced cytotoxicity, consistent with their earlier developmental status and lower antigen exposure.

When we focused specifically on exhausted CD8^+^ T cells, TLN-specific clonotypes were strongly enriched for pre-exhausted states, whereas TM-specific and shared clonotypes were dominated by intermediate exhaustion phenotypes (Figure S6b, c, Table S3). This pattern was preserved across multiple functional metrics, reinforcing that TLN-derived exhausted T cells occupy earlier, more plastic states with higher likelihood of reinvigoration by checkpoint blockade.

Taken together, pseudotime-based grouping of clonotypes revealed a clear developmental ordering across tissues. TLN-specific clonotypes were enriched at early stages, shared clonotypes peaked at intermediate stages, and TM-specific clonotypes predominated at late stages. This stage-resolved structure mirrors the compartmental polarization observed above—lymph nodes supplying newly activated, less differentiated clones, and tumors accumulating chronically stimulated, highly differentiated populations.

### Tumor mucosa enriches germinal center–like B cells and preserves T–B help signaling distinct from metastatic lymph nodes

To characterize B cell diversification across anatomical niches, we analyzed 85,482 B lineage cells and identified 10 transcriptionally distinct subgroups, including naïve (*IGHD*, *S1PR1*), activated (*TNFRSF13B*, *GPR183*), stressed (*DNAJB1*, *DUSP1*) [15], MT-high (*MT-CO1*, *MT-ND6*), CD3⁺ B cells (*CD3D*, *CD3E*), plasma cells (*JCHAIN*, *MZB1*, *IGHG4*), and two germinal center–like populations corresponding to light-zone (GC1) (*RGS13*, *NEIL1*, *BCL6*) and dark-zone (GC2) (*MKI67*, *TOP2A*) phenotypes (Figure 4a, Figure S7a). Major B cell states were proportionally detected across tissues and patients, offering a consistent basis for assessing compartment-specific differences (Figure S7b, c).

**Figure. 4.**
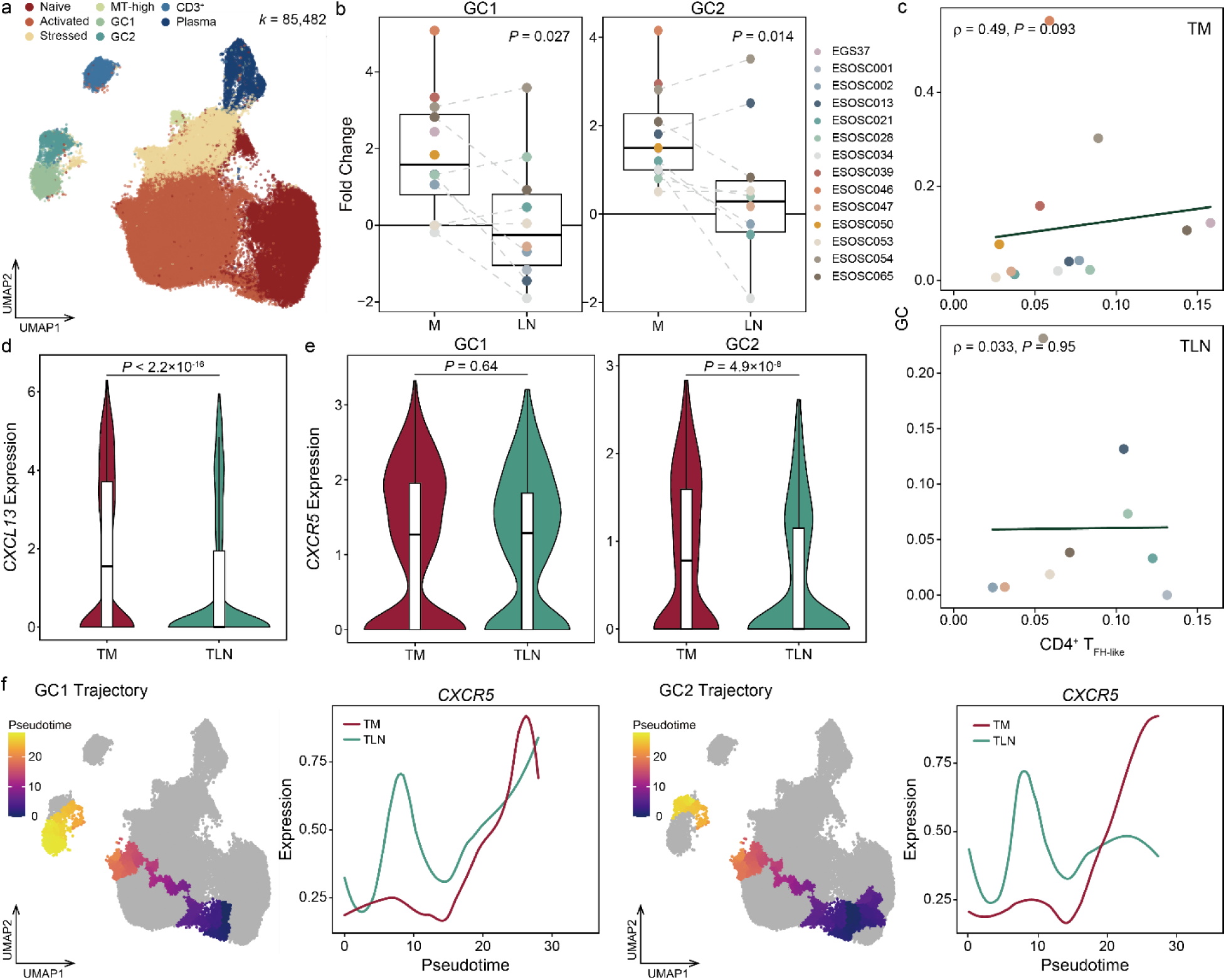
B cell remodeling across ESCC mucosal and lymph node niches. (a) UMAP of 85,482 B and plasma cells partitioned into eight subclusters. (b) Fold changes of GC1 and GC2 B cells fractions across tissue pairs (TM versus NM and TLN versus NLN). Fold changes were calculated per patient (n = 14), and significance assessed using two-sided Wilcoxon rank-sum test. Data are represented as mean ± S.D. (c) Scatter plots showing the relationship between the fraction of GC cells and that of follicular helper–like CD4⁺ T cells (T_FH-like_). Spearman’s correlation coefficient is applied. (d) *CXCL13* expression in T_FH-like_ across TM and TLN indicating ligand availability (two-sided Wilcoxon rank-sum test). (e) *CXCR5* expression in GC B cells across TM and TLN, reflecting receptor readiness for T_FH-like_–B cell interactions (two-sided Wilcoxon rank-sum test). (f) Pseudotime trajectories of GC1 and GC2 subsets and corresponding *CXCR5* expression dynamics along each trajectory.

Focusing on pN^+^ patients, we observed significant enrichment of both GC1 and GC2 subsets in TM relative to TLN (Figure 4b). This expansion suggested that TM sustains a microenvironment permissive to GC-like B cell activation, a finding consistent across patients.

We next examined whether GC B-cell abundance was associated with CD4^+^ T_FH-like_ cells, as these two cell types collaborate in germinal center reactions and TLS. Per-patient correlations showed a modest but TM-specific association between CD4^+^ T_FH-like_ and GC B cell fractions (ρ = 0.49, *P* = 0.093), whereas no relationship was observed in TLN (Figure 4c). This spatially restricted coupling suggested that GC-like B cells and follicular helper-like T cells preferentially co-develop within the primary tumor.

To evaluate whether this enrichment corresponded to functional signaling readiness, we assessed follicular helper-associated and GC-associated transcriptional signatures. CD4^+^ T_FH-like_ cells exhibited significantly higher T_FH-like_ signature scores in TM than TLN (Figure S3d), driven by elevated *CXCL13*, *PDCD1*, *ICOS*, *BCL6*, and *SH2D1A*. In parallel, GC2 cells in TM displayed higher expression of TLS-associated programs and upregulated GC signatures, indicating an activated GC program (Figure S7d, Table S4).

Examining ligand–receptor axes further highlighted compartment-specific tuning of T–B cooperation. In TM, CD4^+^ T_FH-like_ cells expressed significantly higher *CXCL13* (Figure 4d), while GC2 B cells upregulated its cognate receptor *CXCR5* (Figure 4e). Pseudotime analysis revealed two developmental routes, naïve → GC1 and naïve → GC2, both present in TM and TLN (Figure 4f, Figure S7e). However, along the GC2 trajectory—associated with proliferative, DZ-like states—*CXCR5* expression increased preferentially in TM, whereas GC1 trajectories showed comparable dynamics across tissues. These findings suggest that TM may selectively recruit or stabilize GC2-skewed B cell states via a CXCL13–CXCR5 chemotactic axis, rather than globally amplifying GC transcriptional activity.

Together, these observations indicate that primary tumors expand GC-like B cells and maintain strong T_FH-like_–B cell interaction potential, as reflected in increased GC2 skewing, enhanced TLS-associated signatures, and TM-specific CXCL13–CXCR5 signaling. In contrast, metastatic lymph nodes exhibit fewer GC-like B cells and weaker coordination between T_FH-like_ and GC programs, suggesting divergent microenvironmental pressures on humoral immunity across ESCC tissue compartments.

### Compartment-specific myeloid specialization highlights DC activation in TM and *TREM2^high^* macrophage polarization in TLN

We next profiled the myeloid landscape and identified nine transcriptionally distinct subsets among 35,236 cells, including monocytes (*VCAN*, *APOBEC3A*), *TREM2^low^* (*APOE*, *C1QA*, *C1QC*) and *TREM2^high^* macrophages (*TREM2, APOE, C1QA*), *STRAD13⁺* macrophages (*STARD13, CD163, MRC1*), conventional dendritic cells (cDCs) (*CD86*, *CD1C*, *CLEC10*), *CCR7⁺* DCs (*CD86*, *CCR7*), plasmacytoid DCs (pDCs) (*PTCRA*, *CLEC4C*), neutrophils (*FCGR3B*, *S100A8*), and mast cells (*TPSAB1*, *TPSB2*) (Figure 5a). Annotation was supported by canonical lineage markers and functional programs indicative of antigen presentation, migration, or immunoregulatory roles (Figure S8a). The myeloid landscape displayed a broadly preserved architecture across tissues and patients, with all principal subsets detectable in each compartment (Figure S8b, c).

**Figure. 5.**
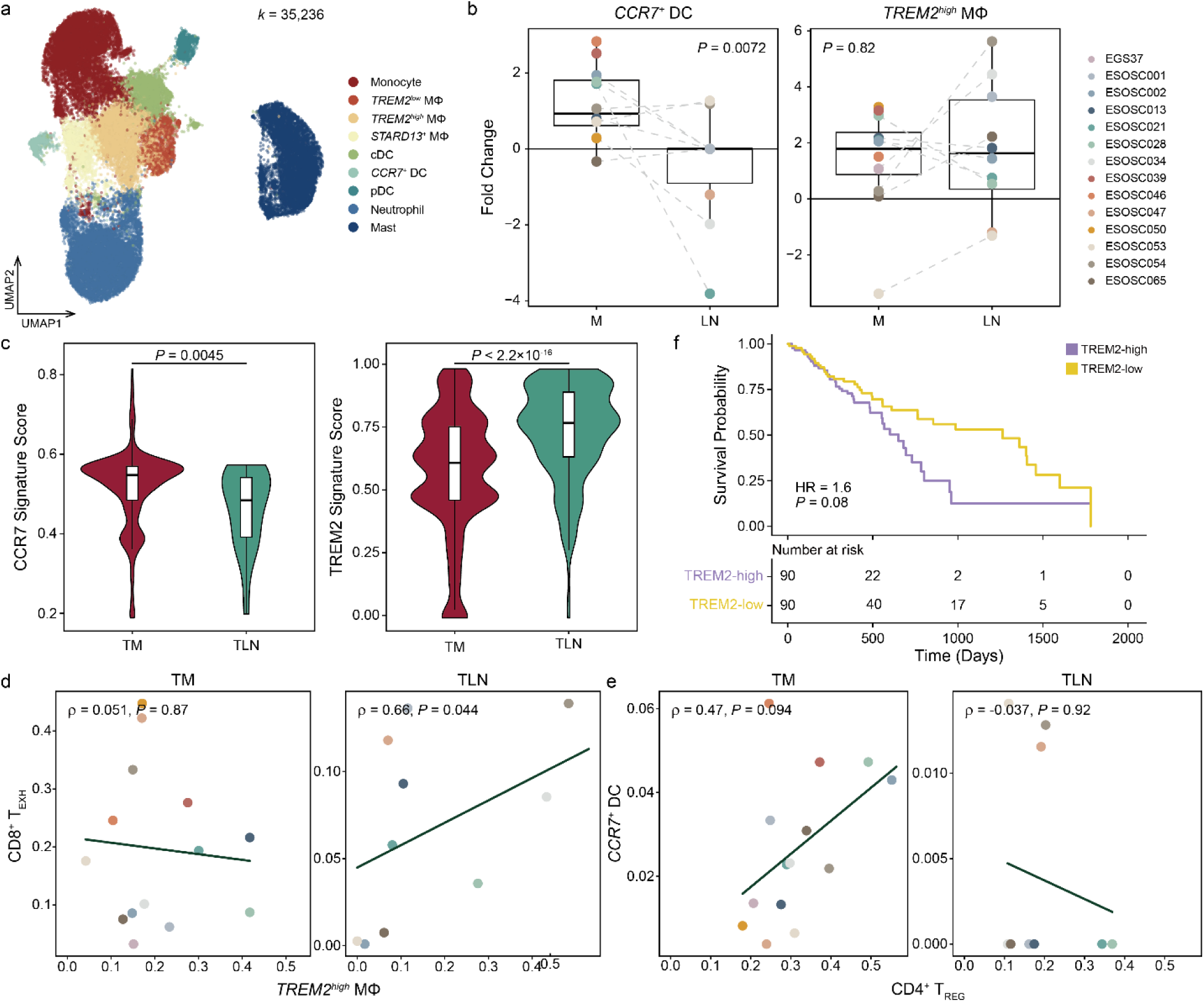
Myeloid diversification and immunoregulatory programs across ESCC sites. (a) UMAP of 35,236 myeloid cells partitioned into nine subclusters. (b) Fold changes of fractions of *CCR7^+^* dendritic cells (DCs) and *TREM2^high^* macrophages between TM versus NM and TLN versus NLN. Fold changes were calculated per patient (n = 14), and significance assessed using two-sided Wilcoxon rank-sum test. Data are represented as mean ± S.D. (c) Signature scores of *CCR7^+^* DCs and *TREM2^high^* macrophages across TM and TLN (two-sided Wilcoxon rank-sum test). (d–e) Scatter plots showing the relationship between the fraction of *TREM2^high^* macrophages and CD8^+^ T_EX_ cells (d), and *CCR7^+^* DCs and CD4^+^ T_REG_ cells (e). Spearman’s correlation coefficient is applied. (f) Kaplan–Meier survival curve based on TREM2-associated signature score. Hazard ratio and *P* value were obtained using multivariable Cox proportional hazards models adjusted for age, sex, and nodal stage.

Across compartments, *CCR7⁺* DCs exhibited a pronounced enrichment in primary tumor relative to metastatic lymph nodes, whereas macrophage populations showed no significant quantitative differences (Figure 5b). *CCR7⁺* DCs have been described as highly activated, migration-competent antigen-presenting cells, and consistent with this phenotype, *CCR7⁺* DCs in TM demonstrated significantly elevated functional scores (*P* = 0.0045), suggesting a qualitatively heightened activation state (Figure 5c, Table S4). In contrast, TREM2 signature scores, which reflects transcriptional programs associated with lipid handling, phagocytosis, and immunosuppression, were markedly higher in TLN than TM (*P* < 2.2 × 10⁻¹⁶). This indicates that although macrophage numbers were similar across niches, *TREM2^high^* macrophages were functionally more polarized in lymph nodes (Figure 5c).

To probe how these myeloid states interface with adaptive immunity, we examined correlations between myeloid subsets and key T-cell phenotypes. In TLN, *TREM2^high^* macrophages showed a positive association with exhausted CD8^+^ T cells (ρ = 0.66, *P* = 0.044) (Figure 5d), whereas in TM, *CCR7⁺* DCs correlated modestly with CD4^+^ T_REG_ abundance (ρ = 0.47, *P* = 0.094) (Figure 5e). These relationships mirror previously reported interactions in which TREM2-driven macrophages reinforce CD8^+^ T cell exhaustion [16], while DC–T_REG_ associations form tolerogenic regulatory circuits that restrain DC activity [17]. Together, these patterns suggest that TLN harbors an immunoactive yet incompletely effective microenvironment in which pre-exhausted CD8^+^ T cells coexist with *TREM2^high^* macrophages that may dampen their potential for reinvigoration, whereas TM shows DC–T_REG_ coupling, reflecting a more fully established immunosuppressive niche.

Given the compartment-specific polarization of *TREM2^high^* macrophages, we assessed the clinical relevance of the TREM2 program using TCGA-ESCA bulk transcriptomes. Patients with higher TREM2 signature expression tended to show significantly poorer survival (Figure 5f). In multivariate Cox models adjusting for age, sex, and pathological nodal stage, nodal status remained an independent prognostic factor, while the TREM2 signature showed a marginal association with outcome (HR = 1.6, CI = 0.95–2.7, *P* = 0.08) (Figure S8d). These results suggest that TREM2-associated macrophage polarization aligns with immunosuppressive features in ESCC, but its prognostic impact may overlap with, or be influenced by, underlying nodal disease burden.

Together, these findings delineate two distinct modes of myeloid regulation across ESCC tissues: (1) an antigen-presenting, *CCR7⁺* DC–dominant state in TM that aligns with regulatory T cell enrichment, and (2) a *TREM2^high^* macrophage–associated immunoregulatory state in TLN that spatially coincides with exhausted CD8^+^ T cells. These dual myeloid programs help explain the divergent immune activation and suppression observed between primary tumors and metastatic nodes and highlight TREM2-associated macrophages as a potential target to enhance CD8^+^ T cell reinvigoration in lymphoid niches.

### Niche-specific fibroblast specialization highlights inflammatory iCAFs in TM and ECM-remodeling CAFs in TLN

To characterize stromal remodeling across ESCC sites, we subclustered 19,052 stromal cells into seven subsets: normal fibroblasts (NF) (*PLA2G2A*, *IGFBP6*), inflammatory cancer-assosicated fibroblasts (iCAFs) (*CXCL1*, *PDGFRA*, *DCN*), myofibroblast-like CAFs (myCAFs) (*COL1A1*, *MFAP2*, *POSTN*), antigen-presenting CAFs (apCAFs) (*CD74*, *HLA-DRA*), *EFNA5⁺* CAFs (*EFNA5*, *DDR2*), smooth muscle cells (SMCs) (*MYH11*, *CNN1*), and pericytes (*RGS5*, *PDGFRB*) (Figure 5a, Figure S9a). While the overall distribution of stromal states across individuals was broadly similar, the marked differences in stromal cell yield between tissues made stable fold-change comparisons less reliable. We therefore assessed directional shifts in abundance rather than applying fold-change metrics (Figure 5b, Figure S9b, c). Both niches demonstrated a decrease in iCAF fractions from normal to tumor, whereas myCAFs expanded substantially in both compartments, with a more pronounced increase in TLN. Notably, apCAFs and pericytes increased in TM but declined in TLN, indicating niche-specific stromal restructuring during tumor progression.

To compare the functional programs of stromal states, we defined CAF modules capturing major fibroblast activities including ECM remodeling, TGFβ signaling, chemokine expression, and antigen presentation (Table S4). This revealed clear compartment-specific specialization (Figure 5c). iCAFs in TM showed significantly elevated chemokine signature scores (*P* = 0.017), consistent with their known role in recruiting immune cells and shaping inflammatory microenvironments in primary tumors. In contrast, both myCAFs and apCAFs exhibited higher ECM remodeling activity in TLN than in TM (*P* = 0.028 and 0.031), suggesting stromal reprogramming as tumor cells establish themselves in secondary lymphoid sites. Antigen presentation scores for apCAFs were modestly higher in TM (*P* = 0.1), aligning with their greater abundance and suggesting a TM-biased antigen presentation program (Figure S9d).

Pathway-level comparison using GSVA refined these observations. In TM, iCAFs exhibited strong enrichment of ECM-related and TGF-β–related programs, including EMT hallmark, ECM degradation, and integrin interactions (Figure 5d, Figure S9e). In TLN, myCAFs showed selective upregulation of ECM organization modules, most prominently within the Hallmark ECM program, reflecting enhanced matrix-structural activity in metastatic nodes. apCAFs, by contrast, demonstrated a mixed pattern: collagen formation pathways were preferentially enriched in TLN, whereas TGFβ hallmark activity remained greater in primary tumor, suggesting that distinct apCAF subsets contribute differentially to stromal programming across compartments.

Together, these data reveal that fibroblast subtypes partition along niche-specific functional axes: iCAFs in primary tumors maintain chemokine-rich inflammatory programs, whereas myCAFs and apCAFs in metastatic lymph nodes exhibit robust ECM remodeling signatures, potentially preparing a permissive microenvironment for metastatic colonization. This division of labor reflects the divergent immunologic and structural demands of primary versus metastatic tumor ecosystems.

### Cell–cell communication mapping reveals distinct immunoregulatory circuits in tumor mucosa and metastatic lymph nodes

To compare microenvironmental communication across niches, we computed average interaction probabilities between major cell types and mapped the differential intensity between TM and TLN (Figure 6a). This analysis revealed broad compartment-specific redistribution of interaction modules, with several signaling axes preferentially enriched in TM and others in TLN (Figure 6b).

**Figure. 6.**
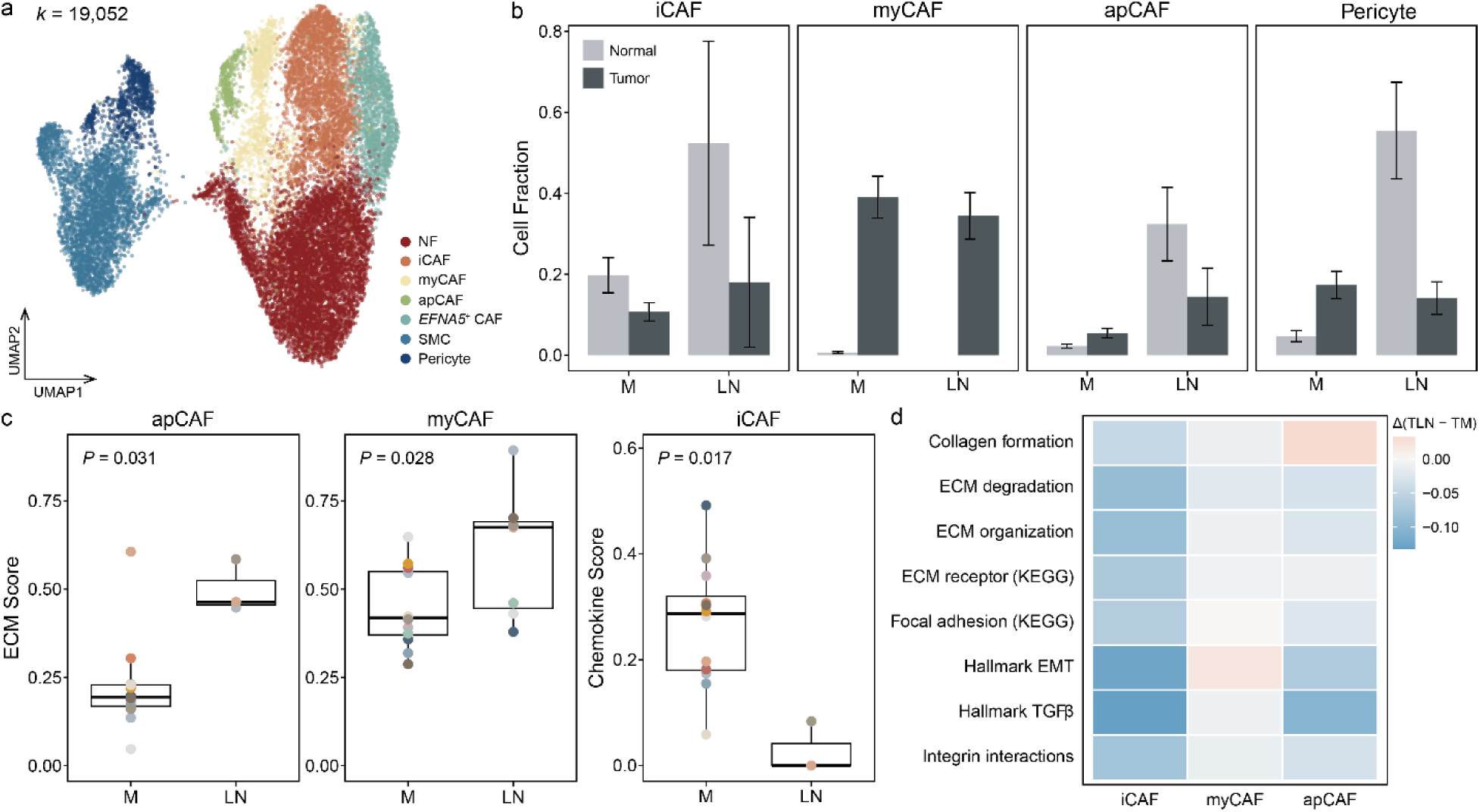
Fibroblast heterogeneity and niche-specific functional programs. (a) UMAP of 19,502 fibroblasts and smooth muscle cells partitioned into seven subclusters. (b) Bar plots show relative cell abundance across compartments. Data are represented as mean ± S.D. (c) CAF-related signature scores across tissue pairs (two-sided Wilcoxon rank-sum test). (d) Heatmap of ECM-associated pathway activity. Values represent Δ(TLN - TM).

TM-enriched interactions were dominated by stroma-derived cues. iCAFs showed the clearest enhancement, displaying stronger communication with CD4^+^ T_REG_ and CD8^+^ T_EXH_ cells, consistent with their chemokine-rich and inflammatory programs in the mucosal niche. Although pericytes and CD4^+^ T_REG_ and CD4^+^ T_FH-like_ cells exhibited mixed TM- and TLN-biased patterns, several TM-skewed interactions emerged repeatedly, including pericyte to *CCR7^+^* DC and T_FH-like_ to GC B cell signaling. These patterns suggest that TM relies heavily on stromal cues to regulate antigen-presenting cell localization and to coordinate local T–B cell coupling.

Pericytes in TM displayed increased CCL19 and CCL21 signaling to *CCR7^+^* DCs (Figure S10a), indicating a stromal positioning mechanism that may enhance the recruitment or retention of migratory DC subsets within the mucosal tumor niche. CD4^+^ T_FH-like_ cells also communicated more strongly with GC2 B cells in TM, echoing earlier findings of coordinated T_FH-like_–GC coupling within the mucosal microenvironment (Figure 4d-e, Figure 6c). Together, these interactions highlight a TM niche structured around chemokine-active fibroblasts and perivascular stromal cells that modulate the abundance and functionality of antigen-presenting DCs. Consistent with this, *CD80/CD86* expression in DCs and *CTLA4* expression in TM-resident T_REG_ cells were significantly elevated, supporting a model in which TM maintains robust antigen presentation while imposing strong T_REG_-mediated regulation (Figure S11d, e).

In contrast, TLN-specific interactions converged around *TREM2^high^* macrophages. These macrophages showed selectively increased SPP1-mediated signaling to CD4^+^ T_REG_ and CD8^+^ T_EXH_ cells (Figure 6d), consistent with their expanded functional signatures in TLN. Supplementary analyses further confirmed significant upregulation of SPP1 in *TREM2^high^* macrophages and corresponding elevation of CD44 and integrin receptors in T_REG_ and T_EXH_ subsets (Figure S11a–c).

These TLN-enriched interactions align with earlier TCR and exhaustion analyses showing that metastatic lymph nodes harbor substantial pools of pre-exhausted CD8^+^ T cells with preserved cytotoxic potential, yet these cells reside within immunoregulatory niches structured around *TREM2^high^* macrophages. The enhancement of *TREM2^high^* MΦ → CD8^+^ T_EXH_ and *TREM2^high^* MΦ → CD4^+^ T_REG_ interactions in TLN suggests that this macrophage program represents a dominant checkpoint limiting full T cell reinvigoration at metastatic sites. Additional interaction pairs, including iCAF → T_REG_ and CXCL12–CXCR4 activity, are shown in Figure S10b.

Together, these findings demonstrate that TM and TLN rely on fundamentally distinct immunoregulatory architectures. TM is shaped by chemokine-producing fibroblasts and activated perivascular stromal cells that orchestrate DC positioning and regulate antigen presentation. TLN instead assembles a focused suppressive network centered on *TREM2^high^* macrophages interacting with exhausted and regulatory T cells. This division of labor explains how metastatic lymph nodes can maintain clonally related, pre-exhausted CD8^+^ T cells with reinvigoration potential while simultaneously imposing strong macrophage-mediated suppression, revealing *TREM2^high^* macrophages as a key therapeutic target within the metastatic niche.

Together, these findings reveal that TM and TLN rely on fundamentally distinct immunoregulatory circuits. TM is shaped by chemokine-producing fibroblasts and pericytes that coordinate DC positioning and regulate antigen presentation through the CD80/CD86–CTLA4 axis of regulatory T cells. TLN instead assembles a *TREM2^high^* macrophage-centered suppressive network targeting exhausted and regulatory T cells. This division of labor explains how metastatic lymph nodes can maintain reinvigoration-competent CD8^+^ T cells while simultaneously imposing macrophage-mediated suppression and identifies *TREM2^high^* macrophages as a promising therapeutic target to enhance immune activation within the metastatic niche.

## Discussion

Our single-cell analysis reveals that the immune microenvironment of esophageal squamous cell carcinoma is highly compartmentalized across primary tumor mucosa and metastatic lymph nodes. These findings provide a mechanistic explanation for long-standing clinical observations that nodal response is a stronger determinant of prognosis than primary tumor regression [18, 19], and that neoadjuvant immunotherapy more effectively eliminates nodal disease than conventional chemoradiotherapy [10]. By mapping 344,790 single cells with matched T cell receptor clonotypes, this study defines the distinct immunologic pressures that shape T cell differentiation, clonal dynamics, and immunosuppressive architecture within each anatomical niche.

A central discovery is that CD8^+^ T cells follow divergent exhaustion trajectories in tumor mucosa and metastatic lymph nodes. Primary tumors were enriched for intermediate and terminally exhausted states. Although intermediate-exhausted cells maintained substantial cytotoxic potential, their elevated inhibitory receptor expression suggested a constrained ability to deploy effector programs, whereas terminally exhausted cells exhibited both high exhaustion and profound loss of cytotoxicity at late pseudotime. In contrast, metastatic lymph nodes contained a greater proportion of pre-exhausted cells that had not yet acquired full exhaustion signatures retaining perforin and granzyme expression. This preservation of effector capacity suggests that nodal CD8^+^ T cells remain in a reversible dysfunctional state, consistent with clinical observations that neoadjuvant chemoimmunotherapy induces robust pathological responses in metastatic lymph nodes [10]. The enrichment of early-state exhausted T cells in lymph nodes aligns with recent notions that tissues differ in their “exhaustion setpoints,” wherein lymphoid niches permit prolonged T cell persistence without irreversible terminal dysfunction. These results provide a cellular basis for the superior nodal downstaging observed with neoadjuvant immunotherapy in ESCC.

Clonal analysis further supports active and dynamic immune surveillance within metastatic nodes. Clonotypes shared between primary and metastatic sites occupied intermediate pseudotime states and exhibited strong cytotoxicity despite exhaustion, indicating that antigen-driven T cell migration maintains functional potential across compartments. Moreover, the correlation between clonal expansion and cytotoxicity was markedly stronger in lymph nodes, whereas in tumor mucosa, clonal growth was more tightly coupled to exhaustion. These patterns imply that proliferating T cells within metastatic nodes remain capable of productive antitumor responses, whereas chronic stimulation in tumors drives cells rapidly toward terminal exhaustion. Together, these results suggest that metastatic lymph nodes serve as a reservoir of rejuvenable tumor-specific T cells and act as a key immunologic site during treatment.

The immunosuppressive mechanisms constraining these T cell responses were also distinct by compartment. In primary tumors, regulatory T cells were quantitatively and qualitatively dominant and engaged in strong CTLA4-mediated suppression of *CCR7^+^* dendritic cells [20, 21]. This interaction likely limits antigen presentation within the tumor bed, contributing to the accumulation of deeply exhausted CD8^+^ T cells. By contrast, metastatic lymph nodes were characterized by a *TREM2^high^* macrophage program that delivered SPP1-dependent suppressive signals to both regulatory and exhausted T cells. These findings align with reports in breast and colorectal cancers that *TREM2^high^* macrophages shape exhausted T cell states and hinder checkpoint blockade [22, 23]. Our study confirms this phenomenon in the context of nodal metastasis and extends it by identifying a spatially organized macrophage centered suppressive circuit unique to metastatic lymph nodes.

Stromal remodeling also diverged across compartments in a manner consistent with their functional roles. Tumor mucosa exhibited chemokine rich inflammatory iCAFs that likely contribute to chronic T cell stimulation and recruitment of regulatory populations. In contrast, metastatic lymph nodes displayed ECM remodeling myCAFs and apCAFs, suggesting matrix restructuring that facilitates metastatic colonization and shapes immune cell positioning. Together with lymph node specific macrophage suppression, this stromal specialization may contribute to the immunologic bottleneck that restrains T cell reinvigoration despite the presence of antigen experienced clones.

These compartment specific insights carry important therapeutic implications (Figure 7e). The dominance of T_REG_ and CTLA4-reliant suppression in primary tumors suggests that CTLA4-based strategies or dendritic cell activation could synergize with PD-1 blockade in the mucosal niche [20]. Our findings further provide mechanistic support for the clinical activity of dual checkpoint blockade in ESCC, in which adding CTLA-4 inhibition to PD-1 blockade has shown improved survival over chemotherapy [24]. The marked accumulation of highly suppressive, CTLA4-expressing T_REG_ in primary tumors, coupled with their direct inhibition of *CCR7^+^* DCs, indicates that CTLA-4 blockade may uniquely relieve a mucosa-specific regulatory bottleneck that PD-1 monotherapy cannot fully overcome. In this model, PD-1 inhibition primarily reinvigorates dysfunctional CD8⁺ T cells, whereas CTLA-4 blockade dismantles the T_REG_–DC axis that constrains antigen presentation and effector priming in the tumor mucosa.

**Figure. 7.**
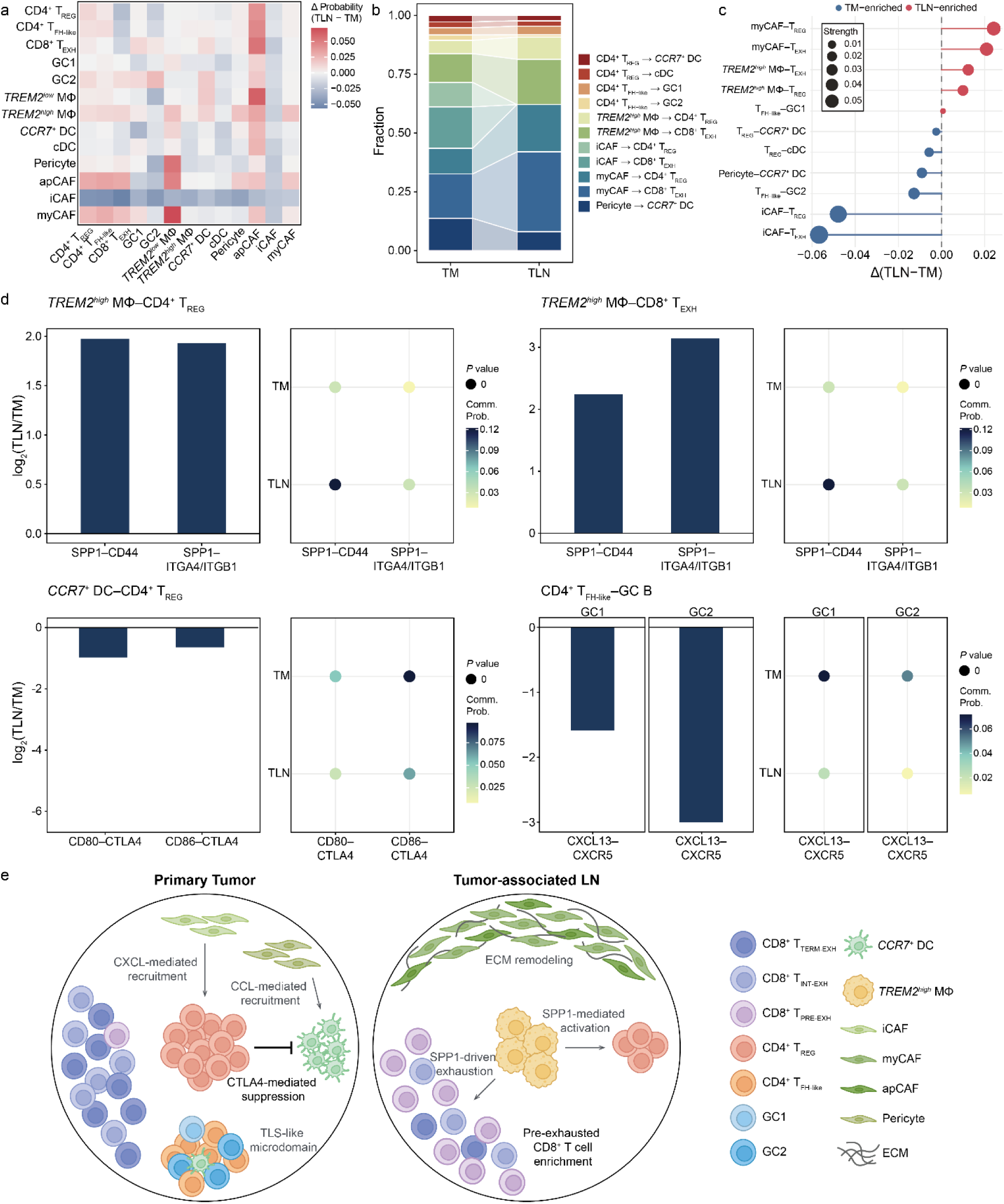
Compartment-specific remodeling of cell–cell communication in TM and TLN. (a) Heatmap showing the average change in interaction probability between TM and TLN, with red indicating higher interaction strength in TLN and blue in TM. (b) Fractional contribution of core interaction pairs in TM and TLN. Each band represents the proportional weight of a specific source–target interaction among the predefined core pairs, illustrating how interaction modules redistribute across compartments. (c) Differential interaction strength for core source–target pairs comparing TLN and TM. Each point represents the signal strength difference (TLN - TM), with color indicating the direction of enrichment and point size reflecting absolute interaction magnitude. (d) Zoomed-in views of representative ligand–receptor pairs illustrating key compartment-specific interaction shifts. Bar plots show log2 fold changes of interaction strength (TLN relative to TM), and dot plots depict communication probability for each ligand–receptor pair in each compartment. (e) Proposed model of the compartment-specific microenvironmental architecture in primary tumors and tumor-associated metastatic lymph nodes of ESCC.

Conversely, the macrophage-centered suppressive axis in metastatic nodes identifies TREM2 inhibition as a promising partner strategy to enhance CD8^+^ T cell reinvigoration [23]. Preclinical investigations of TREM2 blockade have demonstrated potent remodeling of the myeloid landscape. More broadly, these findings underscore the need to evaluate immunotherapy response not only at the primary tumor but also within metastatic lymph nodes, which have historically been overlooked despite being the strongest prognostic determinant in ESCC [18, 19, 25, 26].

Despite the comprehensive nature of this study, several limitations should be acknowledged. First, although the total number of single cells profiled is large, the number of patient samples remains modest due to the rarity of treatment naive, surgically resectable early stage ESCC. This limits the power to fully resolve inter patient heterogeneity. Second, scTCR sequencing was available for a subset of cases, which may underestimate clonal diversity or tissue specific migration patterns. Third, our study focuses on untreated primary and metastatic tissues and therefore cannot directly measure how these compartments remodel in response to immunotherapy. Prospective longitudinal sampling during neoadjuvant treatment will be required to determine whether the pre-exhausted CD8^+^ T cells identified in metastatic nodes truly serve as the cellular substrate of therapeutic response. Finally, functional validation of TREM2-mediated suppression was not performed experimentally, and future organoid or ex vivo coculture systems will be necessary to confirm causality.

In conclusion, this work provides a mechanistic framework explaining why lymph node response is the central determinant of clinical outcome in ESCC and why neoadjuvant immunotherapy preferentially improves nodal clearance [10]. By establishing that tumor mucosa and metastatic lymph nodes harbor distinct exhaustion states, clonal behaviors, stromal programs, and suppressive circuits, this study highlights the necessity of site-specific immunomodulation. Therapies that simultaneously address regulatory T cell suppression in primary tumors and *TREM2^high^* macrophage-driven barriers in metastatic nodes may offer a more complete strategy for durable disease control [23]. Future integration of longitudinal immunotherapy treated samples and functional perturbation studies will help refine these therapeutic opportunities and further clarify how the ESCC immune ecosystem can be reshaped for clinical benefit.

## Methods

### Patient specimens

A total of 18 patients with pathologically confirmed esophageal squamous cell carcinoma (ESCC) were enrolled in this study. Pathological nodal staging comprised four pN0, nine pN1, three pN2, and two pN3 cases. In total, 58 surgically resected specimens were collected, including 19 primary tumor mucosa (TM), 15 adjacent non-tumor mucosa (NM), 14 non-metastatic lymph nodes (NLN), and 10 metastatic lymph nodes (TLN).

All samples were obtained immediately after surgical resection and processed fresh for single-cell analysis. The study was approved by the Institutional Review Board of Yonsei University College of Medicine (IRB Number: 4-2022-0694), and written informed consent was obtained from all participants.

### Tissue dissociation and preparation of single-cell suspensions

Freshly resected tissues were transferred into RPMI-1640 medium (Invitrogen) supplemented with 10% fetal bovine serum and kept on ice. Samples were minced and enzymatically dissociated using the Multi Tissue Dissociation Kit 1 (Miltenyi Biotec) on a gentleMACS Dissociator according to the manufacturer’s instructions. Dissociated cells were filtered through a 70 μm strainer, and erythrocytes were removed using Red Blood Cell Lysis Solution (Miltenyi Biotec). Cell viability and concentration were assessed using a LUNA-FL Automated Fluorescence Cell Counter (Logos Biosystems). Single-cell suspensions were adjusted to the target concentration recommended by 10x Genomics for downstream library preparation. Additional guidance for optimal sample preparation followed the 10x Genomics Single Cell Protocols Cell Preparation Guide (CG000053) and the Guidelines for Optimal Sample Preparation (CG000126).

### Library construction and sequencing

Libraries were prepared using the Chromium controller according to the 10x Chromium Next GEM Single Cell 5’ Reagent Kits v2 and the manufacturer’s user guide (CG000330). Briefly, cell suspensions were diluted in nuclease-free water to achieve a targeted recovery of approximately 10,000 cells, mixed with master mix, and loaded with Single Cell 5′ Gel Beads and Partitioning Oil into a Next GEM Chip K. RNA transcripts from single cells were uniquely barcoded and reverse-transcribed within droplets. cDNA molecules were pooled and enriched with PCR. The amplified cDNA are separated by size selection into 5ʹ Gene Expression and Feature Barcode libraries for TotalSeq-C Hashtag oligos. Feature Barcode (HTO) libraries were enriched using index PCR following the manufacturer’s instructions. For 5’ Gene Expression Library, cDNA underwent end repair, A-tailing, adapter ligation, purification, and subsequent PCR enrichment. Final libraries were quantified by qPCR (KAPA Biosystems) and qualified using the Agilent 4200 TapeStation. Sequencing was performed on an Illumina NovaSeq platform following recommended read configuration.

### Single-cell RNA-seq data processing

Raw BCL files generated by Illumina NovaSeq were converted to FASTQ format using BCL Convert (version 4.3.6). FASTQ files were processed using Cell Ranger (version 8.0.1, 10X Genomics) [27]. Gene expression libraries were analyzed with ‘cellranger count’, and reads were aligned to the GRCh38-2024-A reference genome using the STAR aligner within Cell Ranger. Feature Barcode reads corresponding to TotalSeq-C Hashtag oligos were processed using the Cell Ranger feature-barcode pipeline.

HTO-based demultiplexing was performed in Seurat (version 5.1.0) [28] using the HTODemux function with default parameters, and cells with ambiguous barcode assignment were excluded.

Downstream analyses were performed in R (version 4.4.1). Low-quality cells were removed if they expressed fewer than 200 genes (nFeature < 200) or if more than 20 percent of UMIs mapped to mitochondrial genes (percent.mt > 0.2). Cells with abnormally high UMI counts were removed using a sample-specific threshold defined as the mean plus two standard deviations of total UMIs. Doublets were identified and removed using DoubletFinder (version 2.0.4) [29] with recommendatory parameters. Batch effects among patients were corrected using Harmony (version 1.2.0) [30] prior to clustering and dimensionality reduction.

### Single-cell TCR sequencing data processing

Single-cell V(D)J libraries generated by the 10x Genomics 5′ workflow were processed using Cell Ranger vdj (version 8.0.1) with the GRCh38 reference to assemble TCRα and TCRβ sequences and assign clonotypes. Processed V(D)J outputs were integrated with scRNA-seq metadata using the scRepertoire R package (version 2.2.1) [31]. Cells sharing identical CDR3 amino acid sequences for both TCRα and TCRβ chains were grouped into the same clonotype. Clonal expansion was quantified based on the proportion of cells belonging to each clonotype, and clonotypes were categorized by expansion level using scRepertoire’s default frequency bins. Clonal overlap between tissue compartments was calculated using the Szymkiewicz–Simpson index. TCR metadata were merged with transcriptional clusters to evaluate relationships between clonotype size, exhaustion state, and cytotoxicity signatures.

### Annotation of single-cell RNA-seq data

Following quality control and normalization, scRNA-seq data were analyzed using the Seurat R package. The expression matrix was normalized, and the top 2,000 variable genes were identified using the variance-stabilizing transformation method. Principal component analysis was performed for dimensionality reduction, followed by UMAP for visualization. Cells were clustered using a shared nearest neighbor graph with a resolution of 0.2. Marker genes for each cluster were identified using Seurat’s FindAllMarkers function with the MAST test [32], requiring a log2 fold change greater than 0.25 and expression in at least 25 percent of cells within the cluster. Only positive markers were retained, and *P* values were adjusted for multiple testing. Major cell types were assigned based on canonical marker expression, including T and NK cells (*IL7R*, *CD3D*, *CD3E*, *CD3G*, *TRAC*, *TRBC1*, *TRBC2*, *NKG7*, and *GNLY*), B cells (*MS4A1*, *CD19*, *CD79A*, and *CD79B*), plasma cells (*MZB1*, *JCHAIN*, and *IGHG4*), myeloid cells (*LYZ*, CD14, *CD68*, *CD86*, *MS4A6A*, and *CSF1R*), mast cells (*KIT*, *TPSAB1*, and *TPSB2*), epithelial cells (*EPCAM*, *KRT5*, *KRT7*, *KRT16*, *KRT18*, and *KRT19*), endothelial cells (*VWF*, *PECAM1*, and *EGFL7*), fibroblasts (*DCN*, *C1R*, and *COL1A1*), and smooth muscle cells (*ACTA2*, *MYH11*, *MYL9* and *TAGLN*).

To increase annotation confidence, automated reference-based annotation was performed using Azimuth (version 0.4.6.9001) with the human lung reference atlas (lung_v2, SeuratData) [11]. Each cell received hierarchical prediction scores (levels 1–4), and cells whose Azimuth label matched the manual annotation with a prediction score of at least 0.7 were retained. This concordance-based filtering yielded 344,790 high-confidence cells (91.95 percent of total), which were used for all downstream analyses.

### Cell enrichment analysis

For each patient, the proportion of each cell subtype was calculated within each anatomical compartment (NM, TM, NLN, and TLN) by dividing the number of cells assigned to a given subtype by the total number of cells profiled in that tissue. To enable comparison across patients with different tissue availability, ratios of zero were replaced with a pseudocount of 0.01 before log transformation. Fold change values were computed as log2(TM/NM) for mucosal compartments and log2(TLN/NLN) for lymph node compartments. Wilcoxon rank sum tests were used to assess whether cell-type–specific enrichment differed between mucosa and lymph nodes. All statistical analyses were performed in R.

### Construction of single-cell trajectories

Developmental trajectories were reconstructed using Monocle 3 (version 1.3.7) [33]. For each lineage of interest, annotated Seurat objects were subset to the relevant cell population and converted into Monocle CellDataSet (CDS) objects using the as.cell_data_set function. To ensure pseudotime inference was performed on the same manifold used for cell-type annotation, Harmony-corrected UMAP embeddings and Seurat-derived cluster labels were transferred directly into the CDS object. The principal graph was learned using the learn_graph function with use_partition = FALSE and close_loop = FALSE, yielding a single connected trajectory for each lineage. Root nodes were assigned to naïve or earliest-differentiated clusters based on canonical marker expression, and cells were ordered along the principal graph using order_cells to obtain pseudotime coordinates.

Lineage-specific branches, such as effector versus exhausted paths in CD8+ T cells or GC1-, and GC2-directed branches in B cells, were isolated using the choose_graph_segments function. Pseudotime distributions across tissues were compared using non-parametric tests, and temporal gene expression dynamics were visualized using locally smoothed (loess) regression.

### Cell–cell interaction analysis

Cell–cell communication was inferred using CellChat (version 1.6.1) [34]. For each compartment (TM and TLN), normalized gene expression matrices were used to create CellChat objects. Overexpressed genes and overexpressed ligand–receptor pairs were identified using identifyOverExpressedGenes() and identifyOverExpressedInteraction() under default settings. Communication probabilities were then computed using computeCommunProb() and filtered for robustness using filterCommunication(). Pathway-level communication scores were derived with computeCommunProbPathway(), and aggregated signaling networks were generated using aggregateNet() followed by network centrality analysis via netAnalysis_computeCentrality().

### Gene signature scoring

Gene signature scores were computed using the UCell algorithm (v2.10.1) [35], a rank-based method for single-cell gene set scoring. Log-normalized expression matrices from Seurat objects were supplied to the ScoreSignatures_UCell function together with predefined gene sets, including exhaustion markers (*TOX*, *ENTPD1*, *HAVCR2*, and *PDCD1*) and cytotoxicity genes (*GZMA*, *GZMB*, *PRF1*, *GNLY*, *GZMK*, and *NKG7*). Additional signatures were computed using curated gene sets summarized in Table S4 [16, 36–42]. Resulting UCell scores were stored in the cell metadata and used for comparing states across tissues, cell subsets, and pseudotime trajectories.

### Pathway analysis

Extracellular matrix–associated transcriptional programs were quantified using GSVA (version 2.0.7) with the ssGSEA method [43]. Gene sets corresponding to ECM organization, collagen formation, ECM degradation, integrin signaling, ECM–receptor interaction (KEGG), focal adhesion (KEGG), Hallmark EMT, and Hallmark TGF-β pathways were curated from the MSigDB v2024.1.Hs database via the msigdbr package (version 7.5.1) [44]. For each CAF subtype (iCAF, myCAF, apCAF) in pN+ samples, normalized RNA expression matrices were extracted and converted to uppercase gene symbols to match MSigDB identifiers. Single-sample enrichment scores were computed using ssgseaParam() followed by gsva() with kernel parameters (minSize = 5, α = 0.25, normalization enabled). Resulting pathway scores were aggregated per patient and tissue (TM versus TLN) by averaging cell-level ssGSEA values within each CAF subtype. Differences in ECM program activity between metastatic and primary tumor (TM) compartments were evaluated using paired Wilcoxon signed-rank tests across patient-matched samples, and multiple-testing correction was performed using the Benjamini–Hochberg FDR procedure.

### Survival analysis

Survival analysis was conducted using TCGA-ESCA bulk RNA-seq data and matched clinical metadata [45]. Patients were stratified into high and low groups based on upper and lower quartiles of TREM-associated signature (*TREM2*, *SPP1*, *APOE*, *C1QA*, *C1QB*, and *C1QC*), quantified using UCell scores. Multivariate survival analysis was performed using a Cox proportional hazards regression model from R package survival (version 3.6-4), adjusting for relevant clinical covariates where indicated. Hazard ratios and 95% confidence intervals were computed, and statistical significance was assessed using the Wald test (*P* value < 0.05). Kaplan-Meier survival curves were generated with survminer (v0.4.9), and multivariate model outputs were visualized using forest plots.

### Statistical analysis

Statistical analyses were performed using R (version 4.4.1). Wilcoxon rank-sum test, Wilcoxon signed-rank test, Jonckheere-Terpstra test, and Pearson and Spearman correlation analyses were applied as appropriate. Overall survival was assessed using the Kaplan–Meier estimator with log-rank tests, and multivariable Cox proportional hazards models were used to estimate hazard ratios. Unless specified otherwise, *P* < 0.05 was considered statistically significant.

## Acknowledgement

We thank the patients who participated in this study, as well as their families. This work was supported by a Yonsei-SCL research grant of Yonsei University College of Medicine (6-2024-0152).

