## Supplementary Materials for "Deciphering Tumor Microenvironment Dynamics in Tumorigenesis and Lymph Node Metastasis of Esophageal Squamous Cell Carcinoma using Single-cell RNA Sequencing"

### Supplementary Figures

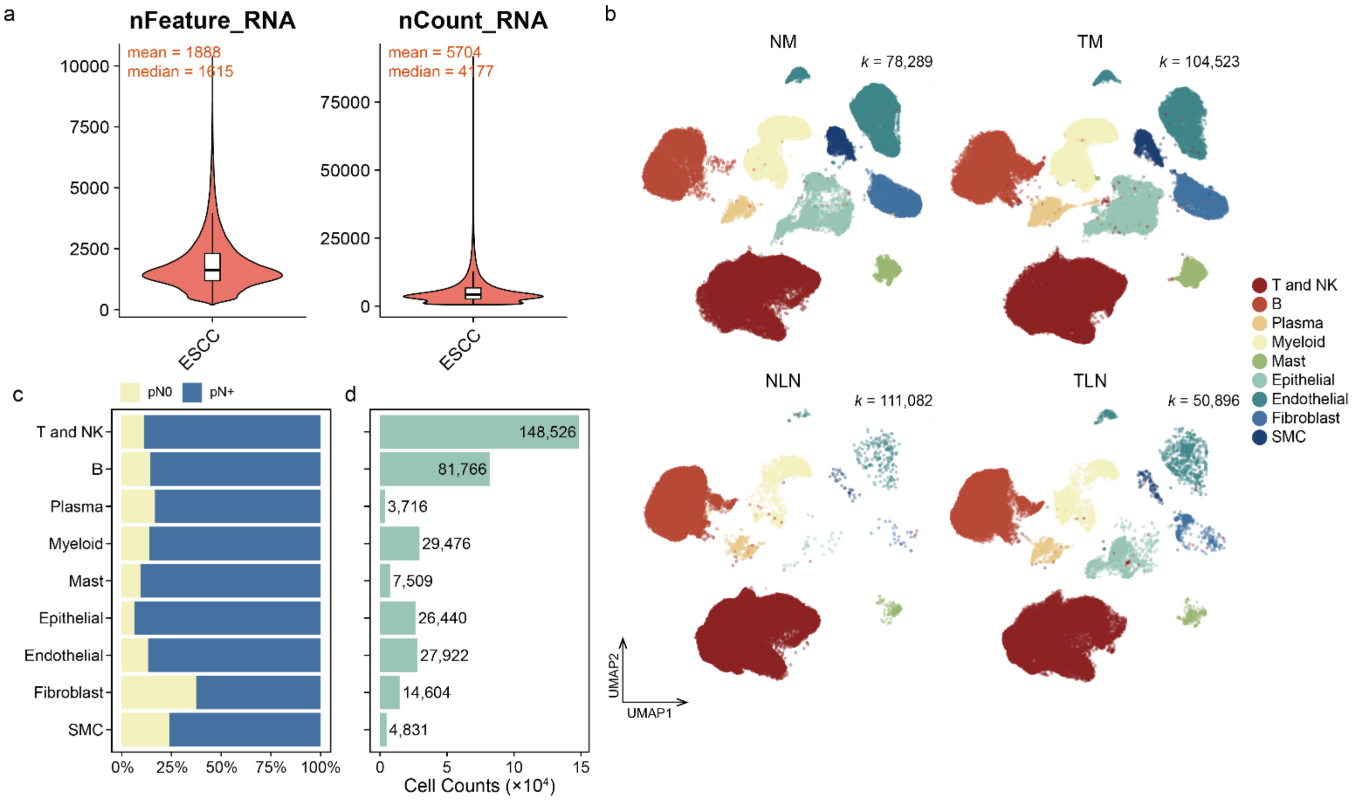


**Figure. S1** | Quality metrics and global cell-type distribution.

(a) Distribution of nFeature and nCount values across all ESCC single cells. Boxes denote median and interquartile range. (b) UMAP embeddings of major cell lineages stratified by tissue compartment (NM, TM, NLN, and TLN), illustrating overall compositional and spatial patterns. (c) Proportion of $pN^0$ and $pN^+$ patients contributing to each major lineage. (d) Total number of cells per lineage aggregated across all tissues.

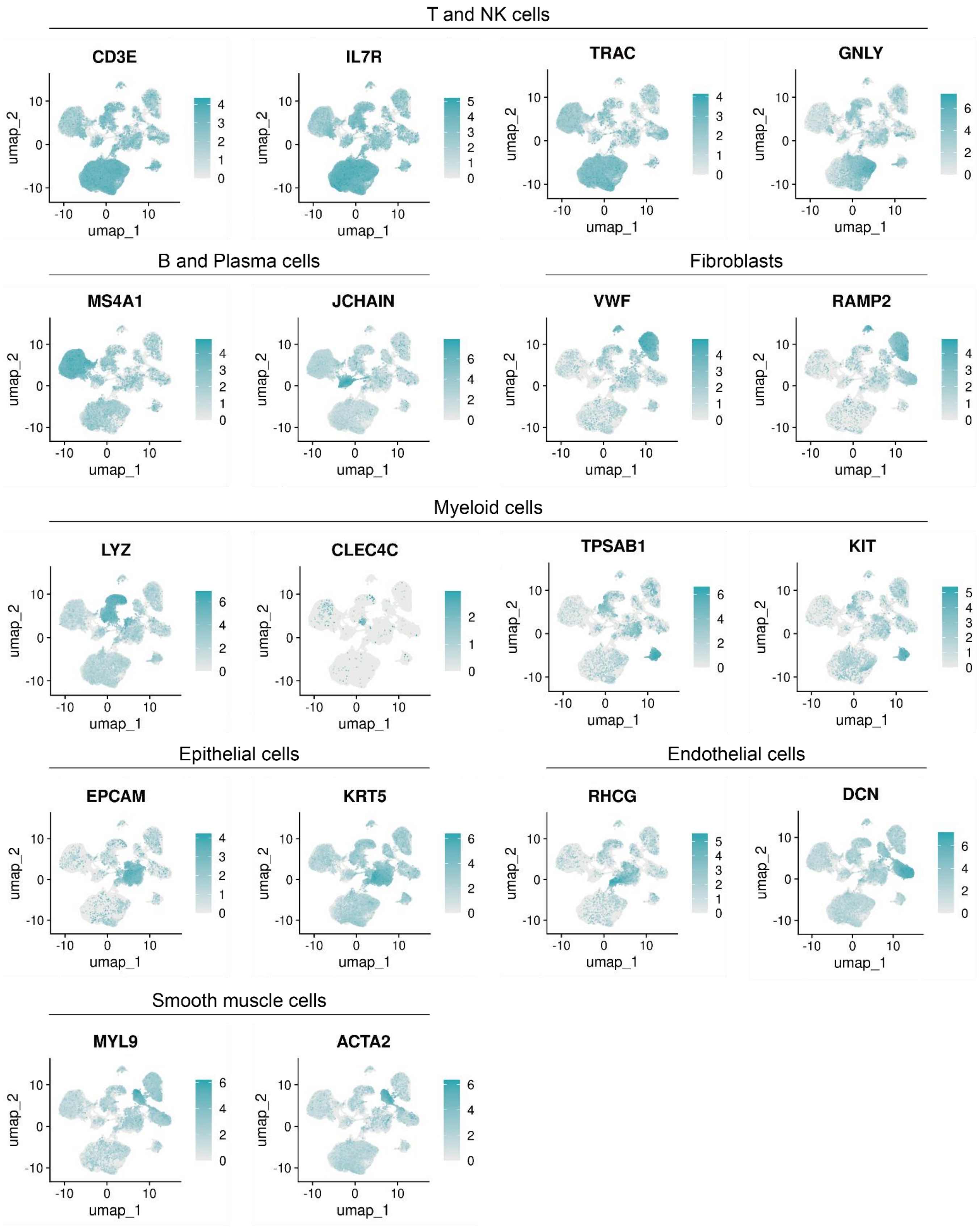


**Figure. S2** | Canonical marker expression across major cell types.

Feature plots showing the expression of representative lineage-defining genes for each major cell compartment. T and NK cells, B cells, plasma cells, myeloid cells, mast cells, epithelial cells, endothelial cells, fibroblasts, and smooth muscle cells are visualized by their canonical markers, confirming robust annotation at the major lineage level.

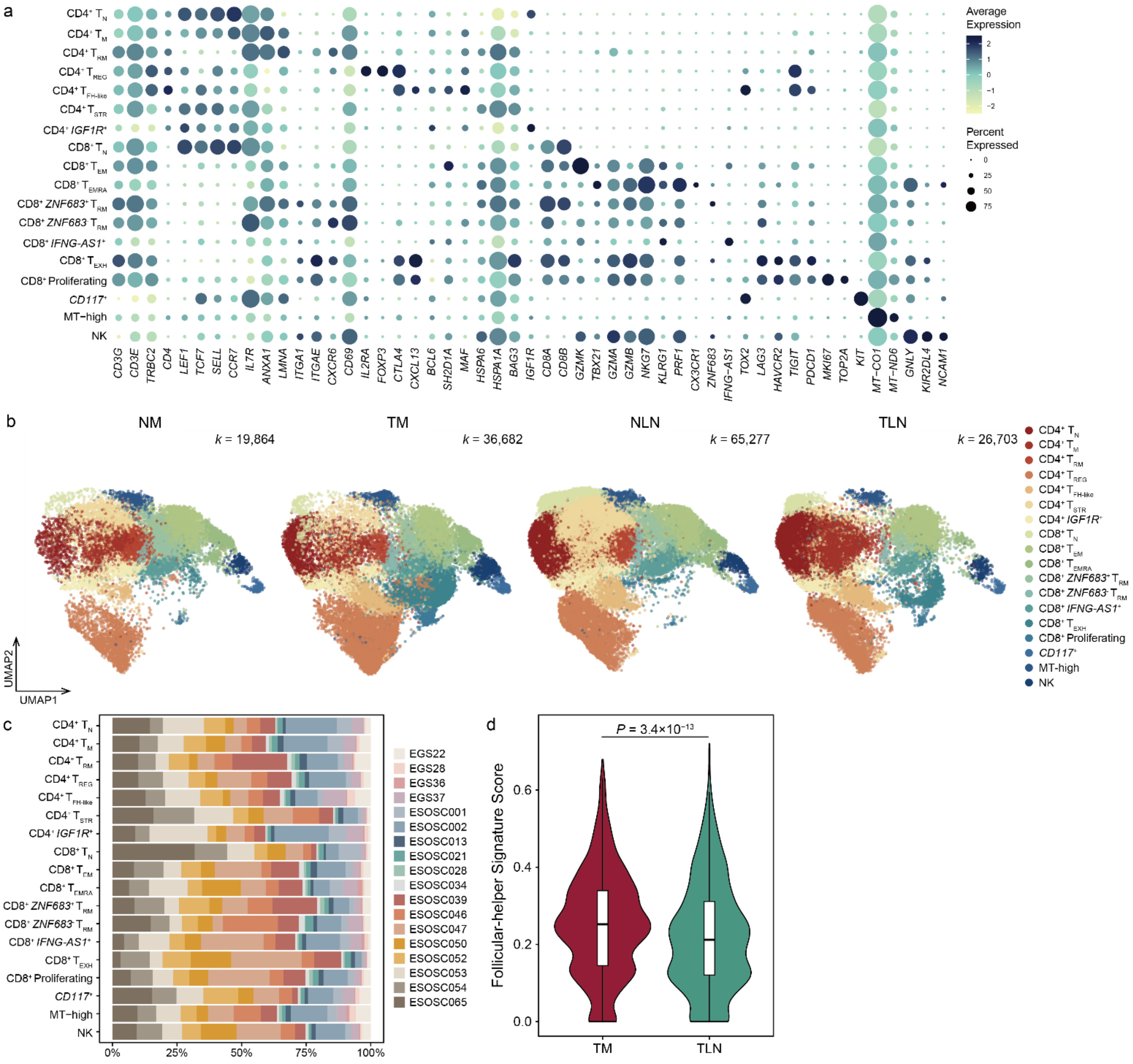


**Figure. S3** | Functional heterogeneity and tissue distribution of T and NK cell subsets.

(a) Dot plot highlighting distinct functional state signatures for each T and NK cell subcluster. Dot size indicates the proportion of cells expressing the marker, while the color intensity reflects the normalized expression level. (b) UMAP of each T and NK cells stratified by tissue compartment. (c) Distribution of patients contributing to each T and NK cell subcluster. (d) Follicular helper signature scores in TM and TLN. *P* value was calculated using the two-sided Wilcoxon rank-sum test.

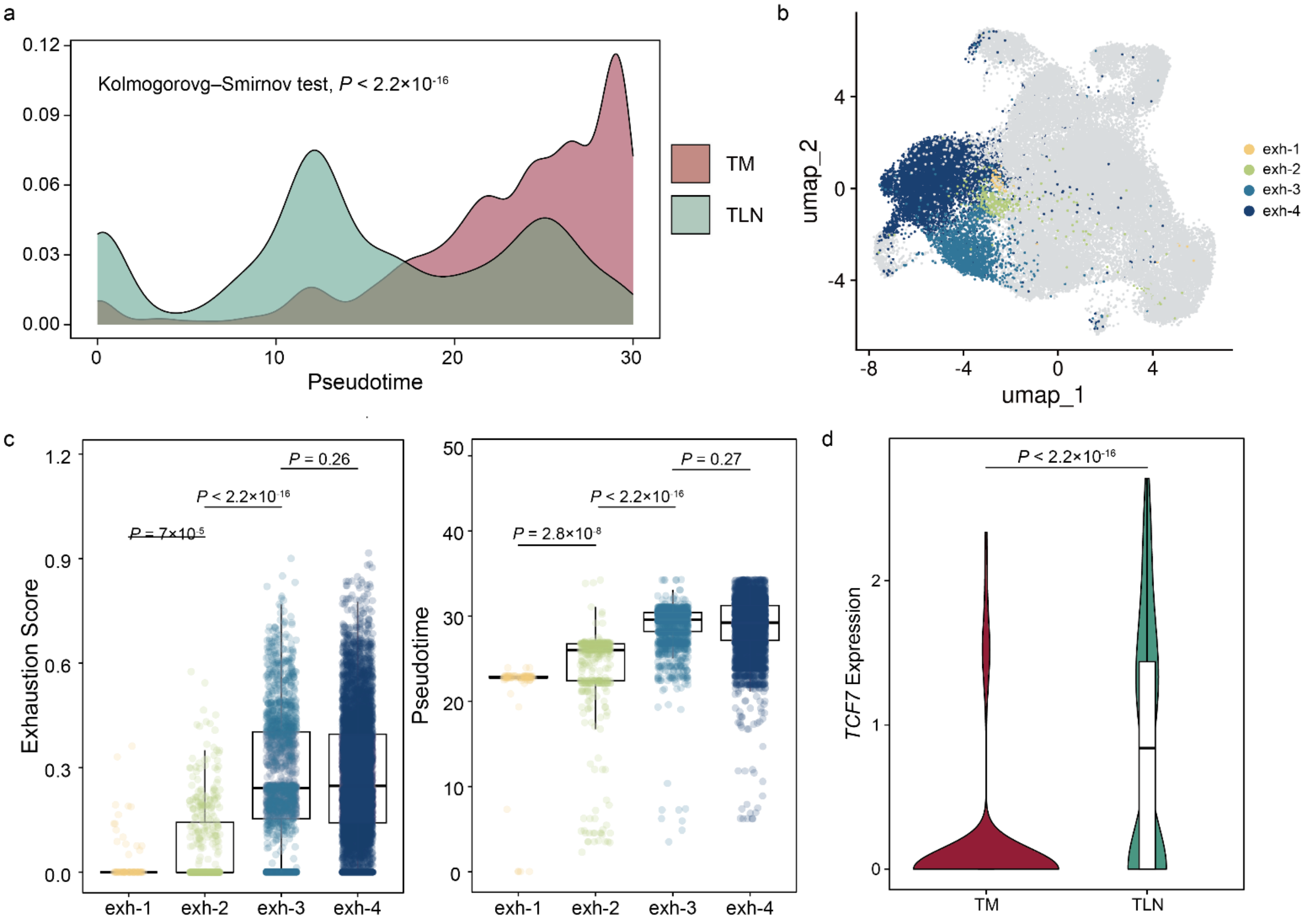


**Figure. S4** | Pseudotime structure and exhaustion-state stratification of CD8$^+$ T cells.

(a) Kernel density estimation of pseudotime distributions for CD8$^+$ T cells in TM and TLN (Kolmogorov–Smirnov test). (b) UMAP visualization of CD8$^+$ $T_{EXH}$ cells partitioned into four unsupervised subclusters. (c) Exhaustion scores (left) and pseudotime values (right) across CD8$^+$ $T_{EXH}$ subclusters. Subclusters exh-1 and exh-2 correspond to early pseudotime ("younger") states, whereas exh-3 and exh-4 represent late pseudotime ("older") states. Exhaustion levels are comparable between the two late subclusters (exh-3 versus exh-4). *P* values were computed using the two-sided Wilcoxon rank-sum test. (d) *TCF7* expression of $T_{PRE\text{-}EXH}$ cells, a two-sided Wilcoxon rank-sum test.

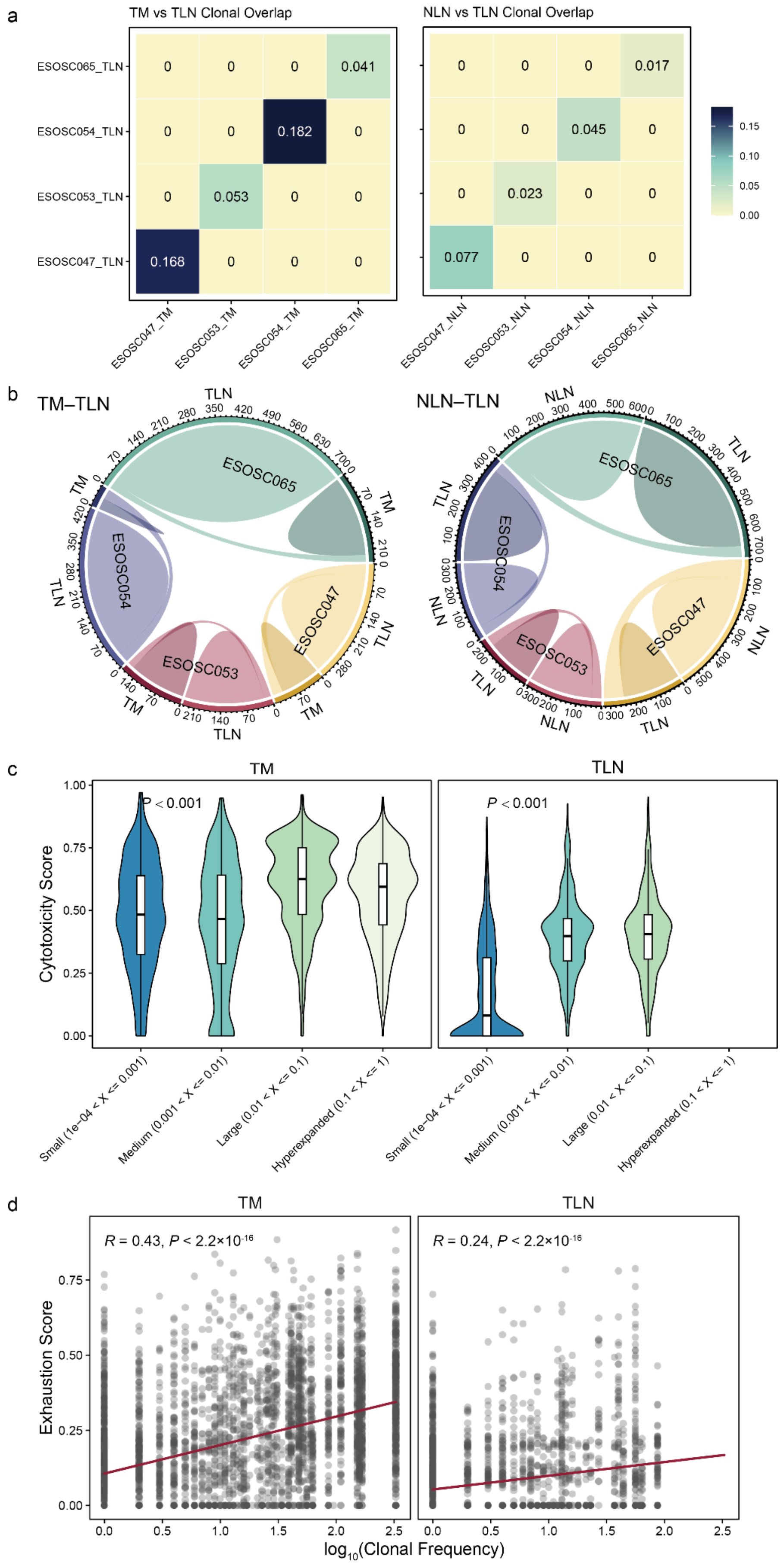
a
TM vs TLN Clonal Overlap
NLN vs TLN Clonal Overlap
ESOSC065_TLN
ESOSC054_TLN
ESOSC053_TLN
ESOSC047_TLN
ESOSC047_TM
ESOSC053_TM
ESOSC054_TM
ESOSC065_TM
ESOSC047_NLN
ESOSC053_NLN
ESOSC054_NLN
ESOSC065_NLN
0.041
0.182
0.053
0.168
0.017
0.045
0.023
0.077
0.15
0.10
0.05
0.00
b
TM–TLN
NLN–TLN
ESOSC065
ESOSC054
ESOSC053
ESOSC047
c
TM
TLN
P < 0.001
P < 0.001
Cytotoxicity Score
Small (1e-04 < X <= 0.001)
Medium (0.001 < X <= 0.01)
Large (0.01 < X <= 0.1)
Hyperexpanded (0.1 < X <= 1)
d
TM
TLN
R = 0.43, P < 2.2×10-16
R = 0.24, P < 2.2×10-16
Exhaustion Score
log10(Clonal Frequency)

**Figure. S5** | Clonal relationships and functional states of CD8$^+$ T cells across tissue niches.

(a) Heatmap of the Szymkiewicz–Simpson overlap coefficient showing clonal sharing between TM and TLN, compared with NLN and TLN, in four patients for whom all required tissue pairs were available. (b) Chord diagrams visualizing the clonal connections between tissue pairs in these four patients, illustrating patient-specific yet reproducible TM–TLN clonal continuity. (c) Cytotoxicity scores across clone-size categories (small, medium, large, and hyperexpanded). Cytotoxic potential increased monotonically with clonal expansion, indicating that expanded clones retain robust effector capacity. Significance was assessed using the one-sided Jonckheere–Terpstra test. (d) Pearson's correlation between clonal frequency and exhaustion of CD8$^+$ T cells.

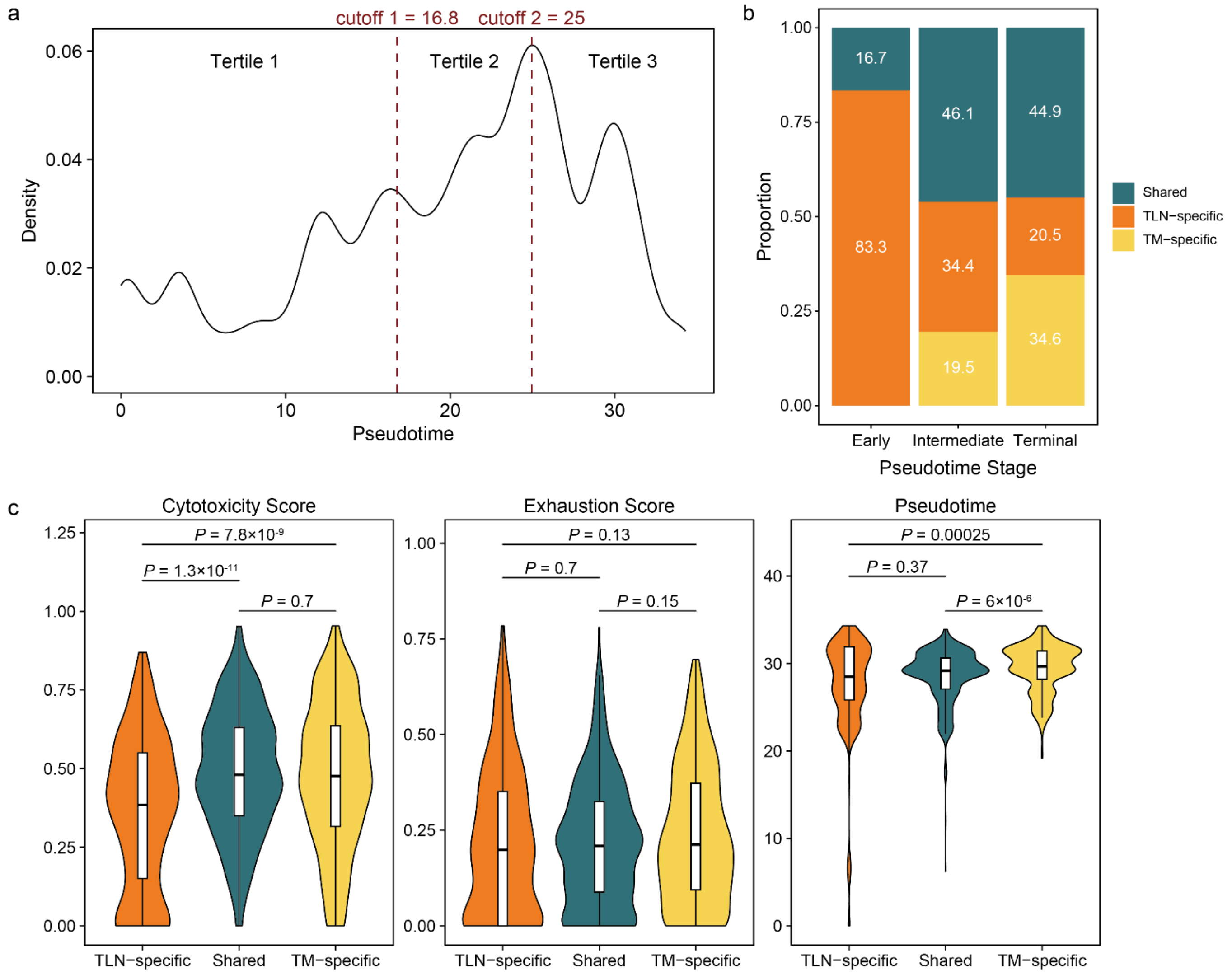


**Figure. S6** | Clonal specialization along the exhaustion continuum of CD8⁺ $T_{EXH}$ cells.

(a) Density plot of pseudotime illustrating the division of the exhaustion continuum into tertiles (early < 16.8, 16.8 ≤ mid < 25, late ≥ 25). (b) Proportional distribution of TM-specific, TLN-specific, and shared CD8⁺ $T_{EXH}$ clones across early, intermediate, and terminal pseudotime stages. (c) Cytotoxicity, exhaustion, and pseudotime scores of CD8⁺ $T_{EXH}$ clones stratified by tissue specificity (two-sided Wilcoxon rank-sum test).

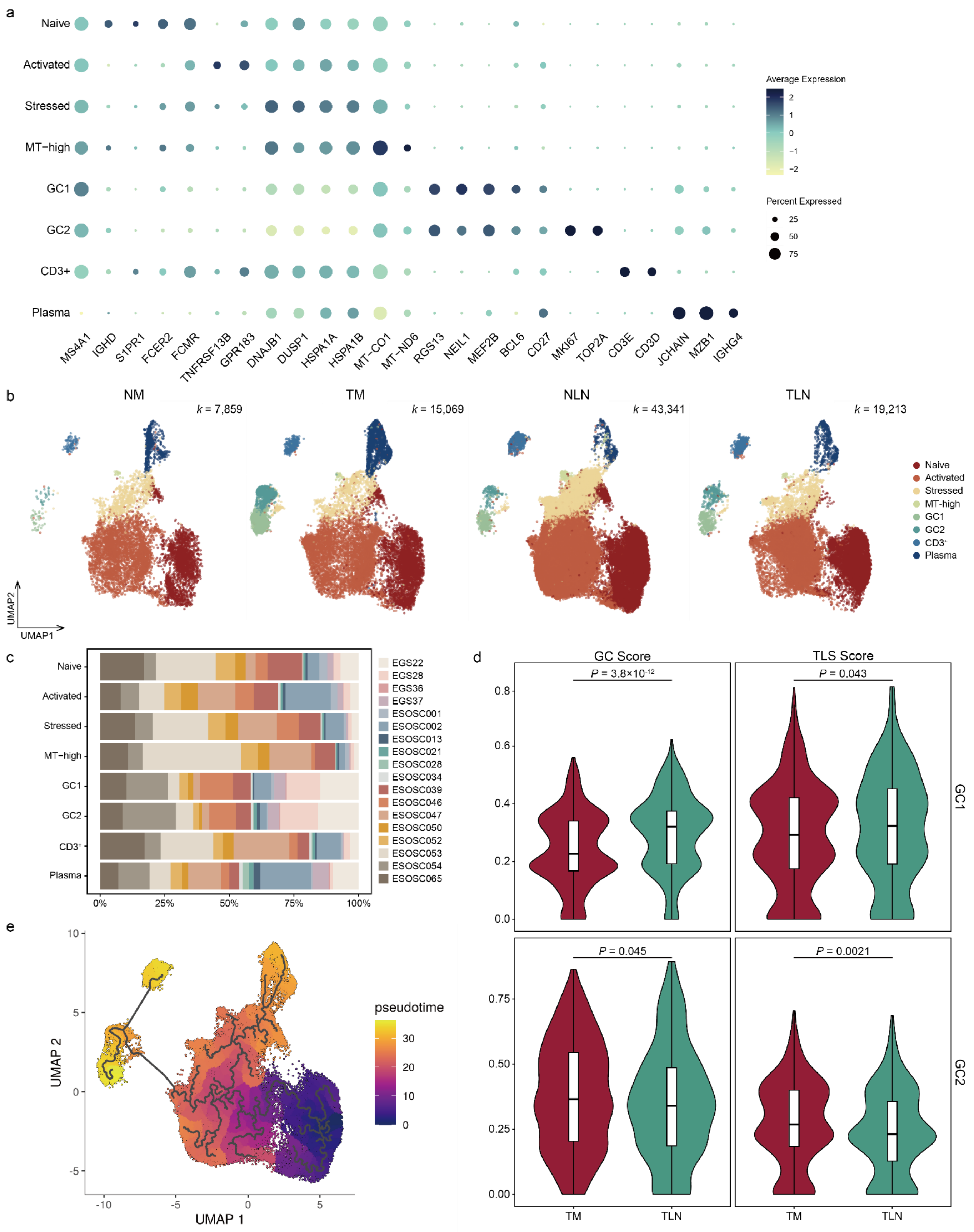


**Figure. S7** | B and plasma cell states across ESCC tissue compartments.

(a) Dot plot showing expression of canonical markers and functional state genes across B and plasma cell subcluster. Dot size indicates the proportion of cells expressing the marker, while the color intensity reflects the normalized expression level. (b) UMAP of each B and plasma cells stratified by tissue compartment. (c) Distribution of patients contributing to each B and plasma cell subcluster. (d) Germinal center (GC) and tertiary lymphoid structure (TLS) signature scores in TM and TLN. Significance was assessed using the two-sided Wilcoxon rank-sum test. (e) Pseudotime projection of B and plasma cells.

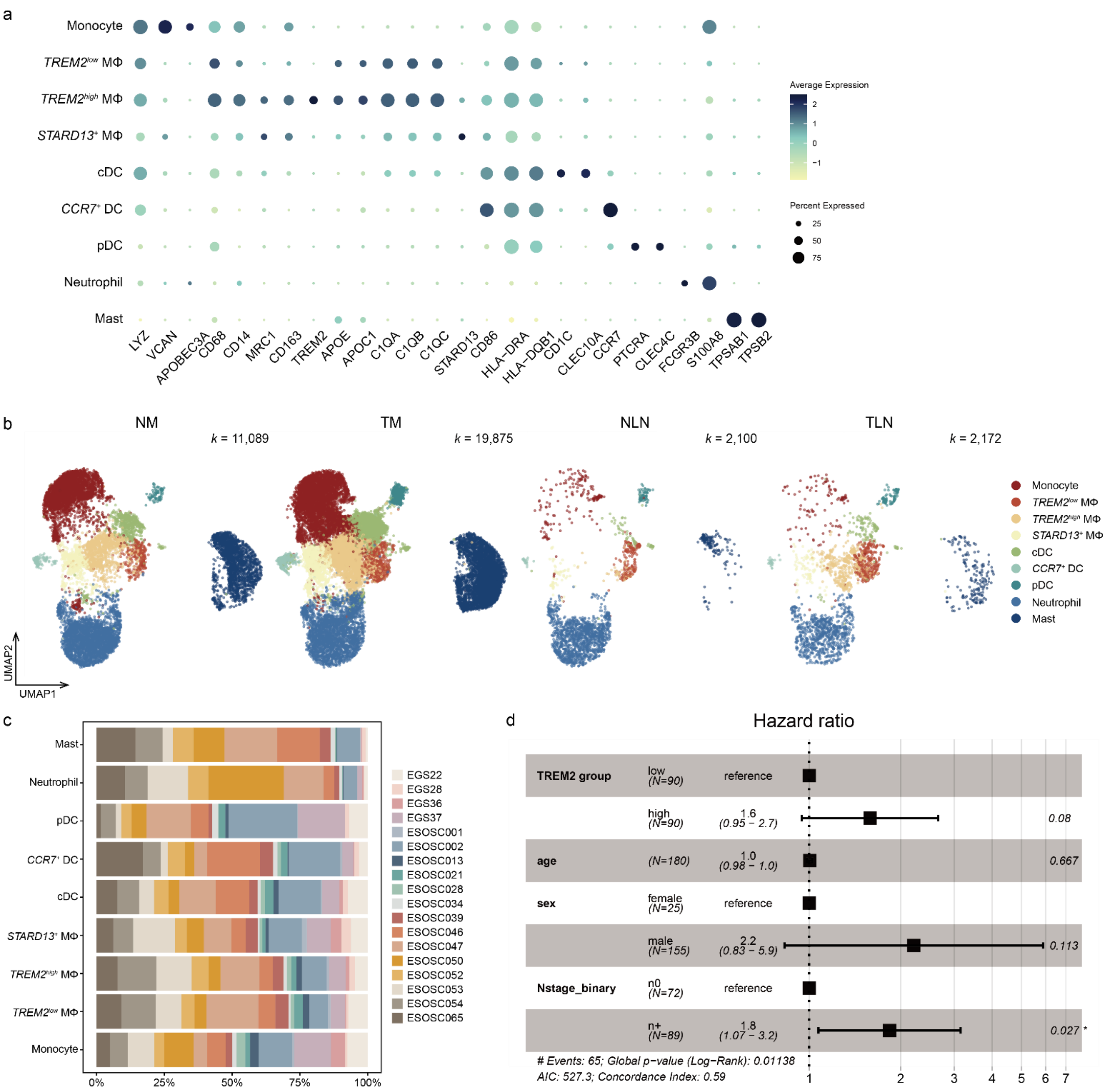


**Figure. S8** | Myeloid and mast cell heterogeneity across tissue niches.

(a) Dot plot highlighting distinct functional state signatures for each myeloid and mast cell subclusters. Dot size indicates the proportion of cells expressing the marker, while the color intensity reflects the normalized expression level. (b) UMAP of each myeloid and mast cells stratified by tissue compartment. (c) Patient distribution across myeloid lineage cells. (d) Forest plot summarizing multivariable Cox proportional hazards analysis evaluating the association between the TREM2-associated macrophage signature (high vs low) and overall survival, adjusted for age, sex, and nodal status. Hazard ratios (HRs) are displayed with 95% confidence intervals (CIs). HRs < 1 indicate reduced mortality risk, whereas HRs > 1 indicate increased risk. Wald test *P* values are shown on the right (**P* < 0.05; ***P* < 0.01; ****P* < 0.001). Group sample sizes are indicated.

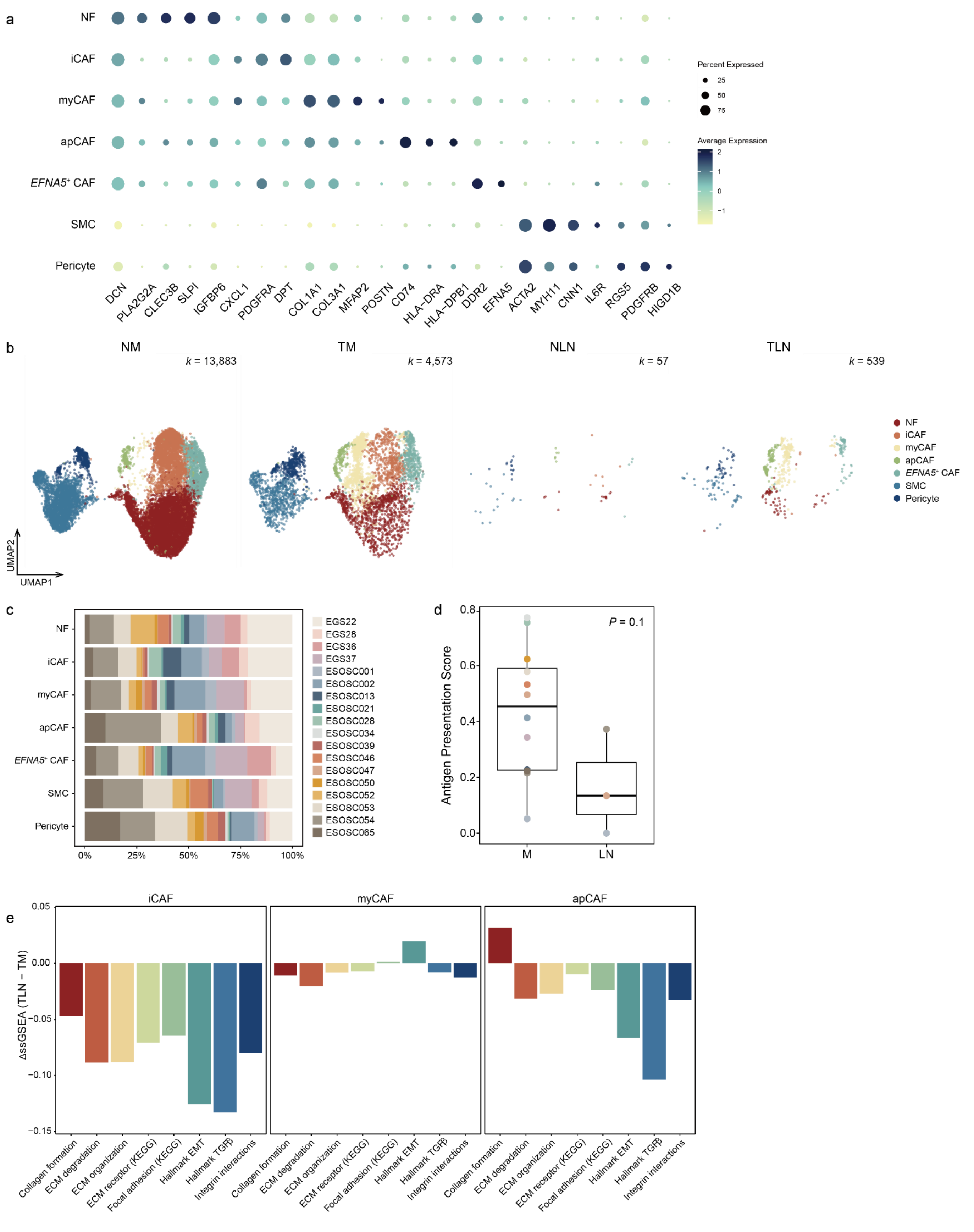


**Figure. S9** | Tissue-dependent remodeling of fibroblast and smooth muscle cell states across ESCC niches.

(a) Dot plot illustrating functional state signatures of fibroblast and smooth muscle cell subclusters. Dot size denotes the proportion of cells expressing each marker, and color intensity reflects normalized expression levels. (b) UMAP of each myeloid and mast cells stratified by tissue compartment. (c) Distribution of patients contributing to stromal cell subcluster. (d) Antigen presentation signature scores of antigen-presenting cancer-associated fibroblasts (apCAFs) across tissue pairs (two-sided Wilcoxon rank-sum test). (e) Bar plots of ECM-associated pathway activity showing $\Delta$(TLN - TM) for each CAF subset.

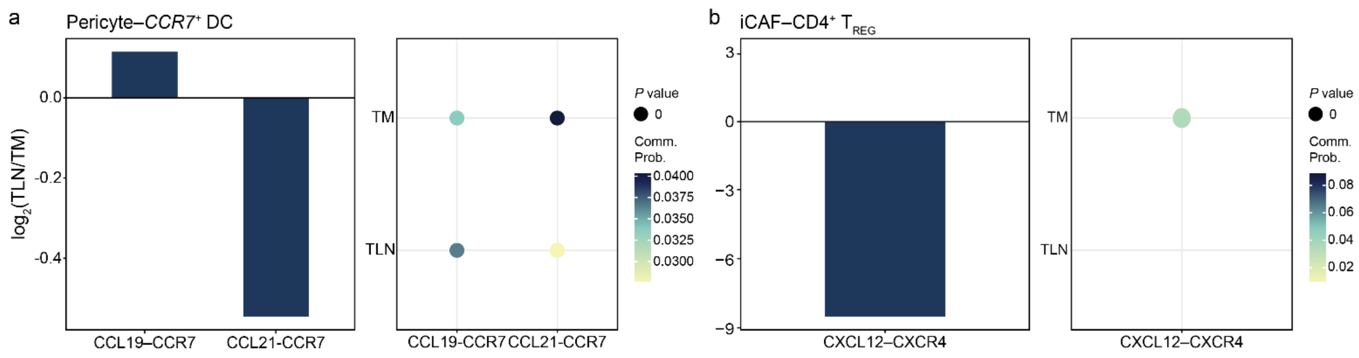


**Figure. S10** | Additional compartment-specific ligand–receptor interaction pairs.

(a-b) Expanded views of ligand–receptor pairs illustrating compartment-specific interaction shifts of pericyte–*CCR7*$^+$ DCs (a) and iCAF–CD4$^+$ T$_{REG}$ (b). Bar plots show log2 fold changes of interaction strength (TLN relative to TM), and dot plots depict communication probability for each ligand–receptor pair in each compartment.

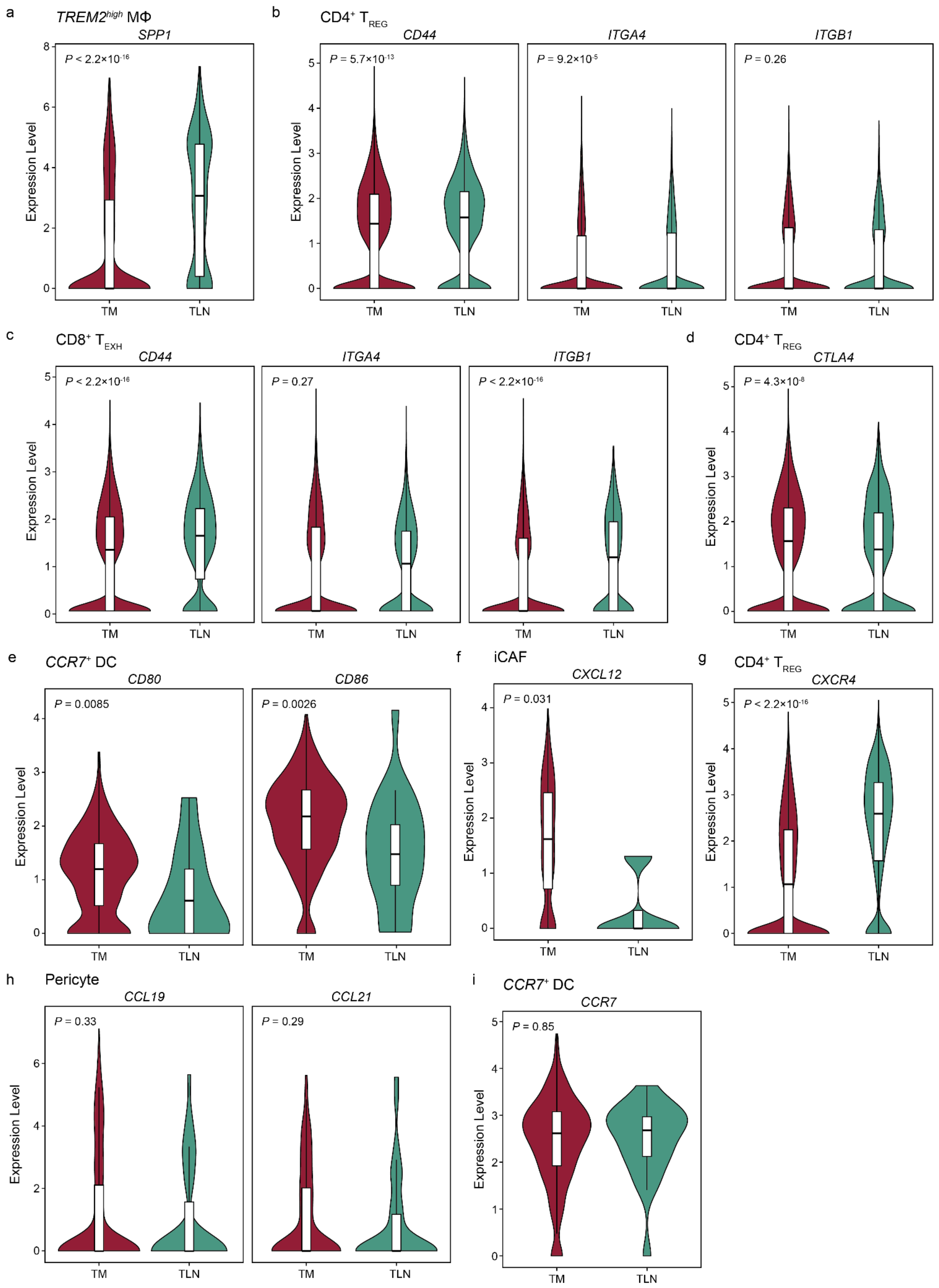
a
TREM2^high MΦ
SPP1
P < 2.2×10^-16
b
CD4+ T_REG
CD44
P = 5.7×10^-13
ITGA4
P = 9.2×10^-5
ITGB1
P = 0.26
c
CD8+ T_EXH
CD44
P < 2.2×10^-16
ITGA4
P = 0.27
ITGB1
P < 2.2×10^-16
d
CD4+ T_REG
CTLA4
P = 4.3×10^-8
e
CCR7+ DC
CD80
P = 0.0085
CD86
P = 0.0026
f
iCAF
CXCL12
P = 0.031
g
CD4+ T_REG
CXCR4
P < 2.2×10^-16
h
Pericyte
CCL19
P = 0.33
CCL21
P = 0.29
i
CCR7+ DC
CCR7
P = 0.85
Expression Level
TM
TLN

**Figure. S11** | Ligand and receptor expression underlying key interaction pathways.

(a-i) Violin plots showing ligand and receptor expression in the relevant source and target cell types across TM and TLN. Each comparison was assessed using a two-sided Wilcoxon rank-sum test.

#### Supplementary Tables

**Table. S1** | Clinical characteristics of patients enrolled in this study.

| Name | Sex | Age | cT stage | cN stage | pT stage | pN stage |
|---|---|---|---|---|---|---|
| Patient 1 | Male | 78 | T1b | N0 | T1b | N0 |
| Patient 2 | Male | 62 | T1b | N0 | T1b | N0 |
| Patient 3 | Male | 67 | T1b | N0 | T1b | N0 |
| Patient 4 | Male | 66 | T2 | N0 | T2 | N1 |
| Patient 5 | Male | 59 | T3 | N1 | T3 | N3 |
| Patient 6 | *NA* | *NA* | *NA* | N1 | *NA* | N1 |
| Patient 7 | Male | 68 | T2 | N1 | T2 | N2 |
| Patient 8 | Male | 62 | T2 | N1 | T1b | N1 |
| Patient 9 | Male | 64 | T2 | N0 | T3 | N3 |
| Patient 10 | Female | 58 | T2 | N0 | T3 | N2 |
| Patient 11 | Male | 59 | T1b | N0 | T1b | N1 |
| Patient 12 | Male | 60 | T1b | N0 | T1b | N1 |
| Patient 13 | Male | 74 | T1b | N0 | T2 | N1 |
| Patient 14 | Male | 58 | T3 | N1 | T3 | N1 |
| Patient 15 | Male | 70 | T1b | N0 | T2 | N0 |
| Patient 16 | Male | 70 | T2 | N0 | T2 | N1 |
| Patient 17 | Male | 64 | T1b | N1 | T1b | N2 |
| Patient 18 | Male | 42 | T1a | N0 | T1b | N1 |

**Table. S2 |** Fisher's exact test results for enrichment of clonotype categories across pseudotime stages.

Fisher's exact tests were performed to evaluate whether TLN-specific, TM-specific, or shared clonotypes were differentially enriched between early, intermediate, and late pseudotime stages. Reported values include raw *P* values, odds ratios, and FDR-adjusted *P* values corresponding to Figure 3e.

| Clonotype | Pseudotime stage comparison | *P* value | Odds ratio | FDR |
|---|---|---|---|---|
| TLN-specific | Early vs. Intermediate | 0.068 | 0.24 | 0.098 |
| | Early vs. Terminal | 0.076 | 0.25 | 0.098 |
| | Intermediate vs. Terminal | 0.85 | 1.05 | 0.85 |
| Shared | Early vs. Intermediate | 0.0013 | 9.39 | 0.003 |
| | Early vs. Terminal | $6.83\times10^{-6}$ | 19.37 | $6.15\times10^{-5}$ |
| | Intermediate vs. Terminal | $6.27\times10^{-4}$ | 2.04 | 0.0019 |
| TM-specific | Early vs. Intermediate | 0.12 | 0.00 | 0.14 |
| | Early vs. Terminal | 0.011 | 0.00 | 0.02 |
| | Intermediate vs. Terminal | $6.10\times10^{-4}$ | 0.46 | 0.0019 |

**Table. S3** | Fisher's exact test results for enrichment of exhausted $CD8^+$ T cell subsets across pseudotime stages.

Fisher's exact tests were performed to quantify whether pre-exhausted, intermediate-exhausted, or terminally exhausted $CD8^+$ T-cell subsets (corresponding to $T_{EXH}$ subclusters in Figure S6b) were differentially enriched across early, intermediate, and late pseudotime stages. Values shown include raw *P* values, odds ratios, and FDR-adjusted *P* values.

| **Clonotype** | **Pseudotime stage comparison** | ***P* value** | **Odds ratio** | **FDR** |
|---|---|---|---|---|
| TLN-specific | Early vs. Intermediate | 0.068 | 0.24 | 0.098 |
| | Early vs. Terminal | 0.076 | 0.25 | 0.098 |
| | Intermediate vs. Terminal | 0.85 | 1.05 | 0.85 |
| Shared | Early vs. Intermediate | 0.0013 | 9.39 | 0.003 |
| | Early vs. Terminal | $6.83\times10^{-6}$ | 19.37 | $6.15\times10^{-5}$ |
| | Intermediate vs. Terminal | $6.27\times10^{-4}$ | 2.04 | 0.0019 |
| TM-specific | Early vs. Intermediate | 0.12 | 0.00 | 0.14 |
| | Early vs. Terminal | 0.011 | 0.00 | 0.02 |
| | Intermediate vs. Terminal | $6.10\times10^{-4}$ | 0.46 | 0.0019 |

**Table. S4** | Gene sets used for UCell signature scoring.

Gene signatures were curated from hallmark pathways, published immune programs, or defined in this study. These gene sets were used for UCell or GSVA scoring throughout the study.

| Signature name | Category | Gene set | Source |
|---|---|---|---|
| Cytotoxicity | T cell | *GZMA*, *GZMB*, *PRF1*, *GNLY*, *GZMK*, and NKG7 | This study |
| Exhaustion | T cell | *TOX*, *ENTPD1*, *HAVCR2*, and *PDCD1* | Bordon, 2019[36]; Gupta et al., 2015[37] |
| Regulatory | T cell | *TGFB1*, *IL10*, *CTLA4*, *LAG3*, *PDCD1*, *CD274*, *ICOS*, *TNFRSF18*, *TNFRSF4*, *FOXP3*, *TIGIT*, *NRP1*, and *IL2RA* | Lucca et al., 2020[38] |
| Follicular-helper | T cell | *CD200*, *CXCL13*, *ICOS*, *PDCD1*, *BCL6*, *SH2D1A*, and *MAF* | Cho et al., 2021[39] |
| Germinal center | B cell | *CXCR4*, *BCL6*, *RGS13*, *MKI67*, and *TOP2A* | Shi et al., 2002[40] |
| Tertiary lymphoid structure | B cell | *CD40*, *CD27*, *SEMA4A*, and *TCL1A* | Ruffin et al., 2021[41]; Xie et al., 2024[42] |
| TREM2 | Myeloid cell | *TREM2*, *SPP1*, *APOE*, *C1QA*, *C1QB*, and *C1QC* | Li et al., 2023[16] |
| CCR7 | Myeloid cell | *CCR7*, *CD86*, *HLA-DRA*, *CD1C*, and *CLEC10A* | This study |
| Chemokine | Fibroblast | *CXCL1*, *CXCL2*, *CXCL8*, *CXCL12*, and *CXCL14* | This study |
| Extracellular matrix | Fibroblast | *ACTA2*, *TAGLN*, *MYL9*, and *COL1A1* | This study |
| Antigen presentation | Fibroblast | *HLA-DRA*, *HLA-DRB1*, and *CD74* | This study |
